# METTL16 promotes taxane resistance in Triple-Negative Breast Cancer through m^6^A-dependent translational upregulation of ABCB1

**DOI:** 10.64898/2026.03.11.710933

**Authors:** Elise G Holvey-Bates, Jesse A Coker, Daniel J Lindner, Anchal Agarwal, Anand Bhushan, Yvonne Parker, Hannah Gilmore, Anton A Komar, George R Stark, Sarmishtha De

**Affiliations:** Department of Inflammation and Immunity, Cleveland Clinic Research, Cleveland, OH, USA; Department of Molecular Medicine, Cleveland Clinic Lerner College of Medicine of Case Western Reserve University, Cleveland, OH, USA; Cleveland Clinic Center for Therapeutics Discovery (C3TD), Cleveland Clinic Research, Cleveland, OH, USA; Department of Shared Laboratory Resources, Cleveland Clinic Research, Cleveland, OH, USA; Center for Gene Regulation in Health and Disease and the Department of Biological, Geological and Environmental Sciences, Cleveland State University, Cleveland, OH, USA; Ophthalmic Research (Cole Eye Institute), Cleveland Clinic, Cleveland, OH, USA; Pathology and Laboratory Medicine Institute, Cleveland Clinic, Cleveland, OH, USA; Department of Biochemistry and Center for RNA Science and Therapeutics, Case Western Reserve University, School of Medicine, Cleveland, OH, USA; Genomic Medicine Institute, Cleveland Clinic, Cleveland, OH, USA

**Keywords:** Triple-negative breast cancer, taxane resistance, METTL16, m^6^A methylation, ABCB1

## Abstract

Triple-negative breast cancer (TNBC) commonly develops resistance to taxane-based chemotherapy, resulting in recurrence and poor clinical outcomes. Defining the molecular mechanisms that sustain chemoresistance is essential for improving therapeutic efficacy. Using unbiased insertional mutagenesis, we identified the RNA methyltransferase METTL16 as a previously unrecognized epi-transcriptomic driver of taxane resistance. METTL16 overexpression conferred resistance to docetaxel and paclitaxel across multiple TNBC models, and METTL16 expression was elevated in paclitaxel-resistant cells. Genetic depletion of METTL16 in paclitaxel-resistant cells restored taxane sensitivity. Because enhanced drug efflux is a well-established mechanism of taxane resistance, we investigated whether METTL16 regulates the multidrug transporter ABCB1 (P-glycoprotein). Paclitaxel-resistant TNBC cells exhibited elevated METTL16 and ABCB1 expression compared to parental cells. METTL16 binds to ABCB1 mRNA and catalyzes its N6-methyladenosine (m^6^A) modification, promoting increased ribosome loading and translational upregulation without altering transcript abundance. Inactivation of METTL16 impaired ABCB1 polysome association and restored paclitaxel sensitivity, demonstrating that the methyltransferase activity is essential for resistance. Consistent with this mechanism, METTL16 overexpression increased ABCB1 protein levels, whereas METTL16 down-regulation increased intracellular paclitaxel accumulation.

Analysis of TNBC patient datasets revealed a positive correlation between METTL16 and ABCB1 expression, supporting the clinical relevance of this mechanism. Antisense-mediated inhibition of METTL16 using a translation-blocking Vivo-Morpholino reduced the survival of resistant TNBC cells and suppressed tumor growth *in vivo*. Surprisingly, genetic ablation of METTL16 caused profound loss of TNBC cell viability, while having only modest effects on nonmalignant mammary epithelial cells, indicating a cancer-selective dependency. Collectively, these findings define a METTL16-ABCB1 interaction that drives taxane resistance and establish METTL16 as a therapeutic target in TNBC.

## Introduction

Advances in systemic chemotherapy have significantly improved outcomes for patients with breast cancer; however, therapeutic resistance limits durable cures, particularly in triple-negative breast cancer (TNBC), which accounts for approximately 15-20% of breast cancers and represents the most aggressive and molecularly heterogeneous subtype (Chen et al. 2025). Because TNBC lacks expression of estrogen receptor (ER), progesterone receptor (PR), and HER2, cytotoxic chemotherapy, especially taxane-based regimens, remains a cornerstone of treatment (Obidiro et al. 2023). However, clinical efficacy is frequently compromised by the rapid emergence of resistance. Therapeutic options for taxane-resistant TNBC are limited, with fewer than 25% of patients responding to alternative treatments, such as platins, poly (ADP-ribose) polymerase (PARP) inhibitors (in BRCA-mutant tumors), or immune checkpoint blockade, and even these responses are typically short-lived (Moreno-Aspitia and Perez 2009; Xu et al. 2024). Elucidating the molecular determinants that enable tumor cells to evade taxane-induced cytotoxicity is therefore critical to improving outcomes in this aggressive disease.

Multiple mechanisms have been implicated in taxane resistance, including activation of pro-survival signaling pathways, alterations in microtubule dynamics, and adaptive changes that limit effective intracellular drug activity (Maloney et al. 2020). We previously demonstrated that overexpression of kinesins, microtubule-based motor proteins, promotes taxane resistance by altering microtubule dynamics (De et al. 2009). Alongside these well-characterized mechanisms, additional regulatory processes contributing to taxane resistance in TNBC remain under active investigation, highlighting the need to identify new therapeutically targetable drivers. In this study, we identified the RNA methyltransferase METTL16 as a previously unrecognized epi-transcriptomic driver of taxane resistance that confers a marked survival advantage under taxane selection.

Regulation of RNA metabolism and function through chemical modification plays a critical role in cancer progression and therapeutic responses (Cerneckis et al. 2024). Formation of N6-methyladenosine (m^6^A), which regulates RNA stability, splicing, translation, and decay, is the most abundant RNA modification (Liu et al. 2017). Among m^6^A writers, METTL16 has emerged as a structurally and functionally distinct, non-canonical methyltransferase that regulates gene expression through both m^6^A-dependent and m^6^A-independent mechanisms (Mermoud 2022; Su et al. 2022). Unlike the canonical METTL3/METTL14 complex which broadly recognizes DRACH (D = A/G/U, R = A/G, H = A/C/U)) consensus motifs, METTL16 exhibits high substrate selectivity, preferentially targeting structured RNA elements that contain the UACAGAGAA/ UACAGARAA motif within stem-loop regions (Pendleton et al. 2017; Satterwhite and Mansfield 2022). This structural specificity enables METTL16 to regulate a limited but biologically critical subset of RNAs, including MAT2A mRNA, U6 snRNA, and the long non-coding RNA MALAT1 (Satterwhite and Mansfield 2022). METTL16 contains an N-terminal methyltransferase domain (MTD) with a Rossmann fold that binds to S-adenosylmethionine (SAM), as well as vertebrate-conserved regions (VCR1/2) that mediate RNA interactions (Satterwhite and Mansfield 2022). The catalytic activity depends on a conserved NPPF motif on the MTD and is further regulated by a unique autoinhibitory K-loop, which modulates SAM accessibility and enzymatic function (Doxtader et al. 2018). These features distinguish METTL16 mechanistically from METTL3/14 and suggest specialized regulatory roles.

METTL16 has been implicated in multiple malignancies, including lung cancer, hepatocellular carcinoma, colorectal cancer, acute myeloid leukemia, ovarian cancer, gastric cancer, pancreatic adenocarcinoma, breast cancer, cholangiocarcinoma, and bladder cancer, where it regulates proliferation, invasion, metabolic adaptation, and immune interactions (Ye et al. 2023; Wang et al. 2025). Despite growing recognition of METTL16 as a selective epi-transcriptomic regulator of cancer-specific phenotypes, its role in resistance to chemotherapy in TNBC is not yet well understood.

Given its established roles in regulating mRNA splicing, stability, and selective translation, METTL16 is positioned to directly control molecular pathways that enable tumor cells to survive and withstand cytotoxic chemotherapy. One of the most clinically relevant mechanisms of taxane resistance is enhanced drug efflux, mediated by ATP-binding cassette (ABC) transporters. Among these, ABCB1 (P-glycoprotein) is a central mediator of multidrug resistance that actively exports taxanes from cancer cells, thereby reducing intracellular drug accumulation and cytotoxicity (Abd El-Aziz et al. 2021). Elevated ABCB1 expression has been strongly associated with chemotherapy failure and poor prognosis in breast cancer patients, yet direct pharmacological inhibition of ABCB1 has been unsuccessful due to the high toxicity of inhibitors and the development of resistance mechanisms (Abd El-Aziz et al. 2021).

We now show that METTL16 binds directly to ABCB1 mRNA and enhances its m^6^A methylation in taxane-resistant TNBC cells, leading to increased ABCB1 expression and drug efflux. These findings position METTL16 upstream of a central multidrug resistance effector, indicating that epi-transcriptomic regulation, rather than transporter blockade, may be more effective in suppressing ABCB1-driven chemoresistance.

To explore therapeutic strategies for targeting METTL16, we utilized antisense approaches that directly inhibit its expression at the translational level. Antisense morpholino oligonucleotides are chemically modified nucleic acid analogs that bind complementary RNA sequences and sterically block processes such as translation initiation without inducing RNA degradation (Moulton and Jiang 2009). Unlike many antisense platforms, morpholinos exhibit high sequence specificity, strong nuclease resistance, and minimal off-target effects, ensuring efficient delivery and sustained activity *in vivo* (Ferguson et al. 2013; Pucci et al. 2024). Importantly, morpholino-based therapeutics have demonstrated favorable safety profiles in both preclinical and clinical settings with FDA-approved applications (Aartsma-Rus and Krieg 2017; Roshmi and Yokota 2023), supporting their translational potential. These properties make morpholino-mediated inhibition a promising strategy for selectively targeting METTL16 and evaluating its therapeutic potential in chemo resistant TNBC.

## Materials and Methods

### Cells and Reagents

MDA-MB-231, MDA-MB-468, and A549 cells were purchased from ATCC and were grown in RPMI 1640 medium. SUM159PT and HCC70 cells, kindly provided by Dr. Ruth Keri (Cleveland Clinic, Cleveland, OH), were grown in DMEM-F12 or RPMI 1640, respectively. The MDA-MB-231-PacR and SUM159PT-PacR cells, also a kind gift from Dr. Ruth Keri, were maintained in RPMI 1640 or DMEM-F12, respectively, supplemented with paclitaxel (100 nM) for three days per week. The media for all these cell lines were supplemented with 5% (vol/vol) heat-inactivated FBS (Biowest), 100 units/mL penicillin, and 100 μg/mL streptomycin. hTERT-HME1 cells were grown in MEGM Mammary Epithelial Cell Growth Medium BulletKit from Lonza (cat. # CC-3150). All cultures were maintained at 37°C in a humidified incubator at 5% CO_2_ and were periodically checked for mycoplasma using Lonza’s MycoAlert Mycoplasma Detection Kit (cat. # LT07-218). Cell lines were carried for no more than 10 passages. The antibody against METTL16 was from Cell Signaling (cat. # 87538S), and that against ABCB1 was from Proteintech Group (cat. # 22336-1-AP). GAPDH-HRP antibody was from Proteintech (cat. # HRP-60004). Anti-β-actin was from Sigma Aldrich (cat. # A5316). Secondary antibodies HRP-goat anti-rabbit IgG (cat. # AS014) and HRP-goat anti-mouse IgG (cat. # AS003) are from Abclonal. Anti-FLAG® M2 antibody (cat. # F1804-50ug) was from Sigma Aldrich. Docetaxel (cat. # S1148) and Paclitaxel (cat. # S1150) were from Selleck Chemicals. For *in vitro* studies, drugs were formulated in 100% DMSO to make 10 mmol/L stock solutions and aliquoted for long-term storage at −20 °C. Transfection reagents Lipofectamine (cat. # 18324012) and PLUS Reagent (cat. # 11514015) were purchased from Invitrogen. Puromycin was from Sigma Aldrich (cat. # p7255-25mg) and hygromycin B was from Invitrogen (cat. # 10687010).

### Constructs, Virus Production, and Infection

Validation-based insertional mutagenesis (VBIM) vector constructs and their use were described previously (Lu et al. 2009; De et al. 2009; Tan et al. 2012). To generate WT METTL16 expression constructs, total RNA was isolated from human cells using a Qiagen RNeasy Mini kit (cat. # 74104) according to the protocol provided by the manufacturer. Reverse transcription was performed with the SuperScript III First-Strand Synthesis System kit and protocol (Invitrogen, cat. # 18080051). Standard PCR reactions were performed using METTL16-specific forward and reverse primers with a stop codon in the reverse primer. The PCR product was cloned into lentiviral vector pLCMV-puro at Age1 and Sal1. The N184A METTL16 mutant construct was generated by site directed mutagenesis. To generate FLAG-tagged constructs, WT and N184A METTL16 were PCR amplified from the pLCMV-puro constructs using a 5’ primer with an XhoI site and a 3’ primer with an XbaI site. PCR products were cloned into the XhoI/XbaI sites of pENTR4-FLAG (Addgene # 17423), in frame with the FLAG tag (N-terminus). Gateway recombinational cloning was performed to transfer FLAG-tagged WT or N184A METTL16 to the pLenti-CMV-puro destination vector (Addgene # 17452). shRNAs against human METTL16 (TRCN0000134067, TRCN0000136815, and TRCN0000136881) were obtained from Sigma-Aldrich (in pLKO.1 vector). lentiCRISPRv2 (Addgene # 52961) was used to generate constitutive Cas9 expressing cells. Lentiviral vectors (lenti-sgRNA hygro, Addgene # 104991) containing METTL16-targeting sgRNA (sgME) or control sgRNA (sgNS) were kindly provided by Drs. Rui Su and Jianjun Chen (Beckman Research Institute of City of Hope, Monrovia, CA).

To produce infectious lentivirus, HEK-293T cells (4 x 10^6^ cells on a 10 cm plate) were transiently transfected with 1.6 μg pCMVR8.74 (Addgene # 22036) and 1 μg pMD2.G (Addgene # 12259) helper plasmids as well as 3 μg lentiviral target vector using Lipofectamine (30 μl) and PLUS Reagent (20 μl) in 6 mL of optiMEM. The virus produced was collected 24 and 48 h after infection, 4 μg/mL of polybrene was added, and the preparation was used to infect human cancer cell lines (MDA-MB-231, MDA-MB-468, SUM159PT, HCC70) and hTERT-HME1 cells.

### Knocking out METTL16 genes using CRISPR-Cas9

Pools of MDA-MB-231, SUM159PT, and hTERT-HME1 cells constitutively expressing Cas9 were generated by infection with lentiCRISPRv2 lentivirus and, 72 h after infection, cells were selected with 3 μg/mL puromycin for the TNBC cell lines and 1 μg/mL for HME cells. Puromycin selection was completed in 3 days. The constitutive-Cas9 expressing cells were then infected in 10 cm plates with lenti-sgNS hygro and lenti-sgME hygro lentivirus. Prior to selection with hygromycin, cells were plated onto 6-well plates for crystal violet staining, 6 cm or 10 cm plates for collecting protein, and 48-well (HME) or 96-well (TNBC lines) plates for cell survival assays. TNBC cell lines were treated with 200 μg/mL hygromycin while HME was treated with 100 μg/mL. Crystal violet staining was performed when sgNS cells reached confluence: 5 days hygromycin treatment for HME and 11 days for MDA-MB-231. Cell lysates were prepared on the same days as the crystal violet assay for HME and MDA-MB-231, and on day 5 of hygromycin treatment for SUM159PT. Cell survival assays were performed after 13 days of hygromycin treatment for MDA-MB-231 (4,000 cells per well), 7 days for SUM159PT (500 cells per well), and 13 days for HME (10,000 cells per well).

### Cell survival assay

Paclitaxel-resistant cells were cultured without the maintenance dose of paclitaxel for at least 5 days prior to experiments. Cells (20,000 per well) plated in 24-well plates were allowed to attach overnight and then treated with docetaxel or paclitaxel for 24 h. After 5 days, cells were lysed with 1 mol/L NaOH, and lysates were diluted 50-fold prior to measuring absorbance at 260 nm as an indicator of total nucleic acid content. The fraction of surviving cells was calculated relative to untreated controls (De et al. 2009). For some experiments, 10,000 cells were plated per well in 96-well plates, allowed to attach overnight, and treated with paclitaxel for 24 h. After 3-5 days of treatment, cell survival was determined using the CyQUANT Direct Cell Proliferation Assay (Thermo Fisher Scientific, cat. # C35011) according to the manufacturer’s instructions, as previously described (De et al. 2018).

For CRISPR-Cas9-mediated METTL16 knockdown cells, cell survival was assessed by crystal violet staining or by using the Alamar Blue reagent (Thermo Fisher Scientific, cat. # DAL1025) according to the manufacturer’s instructions. Briefly, Alamar Blue reagent was added directly to the culture medium in 96-well plates. After incubation for 2.5 h at 37°C, fluorescence was measured using a plate reader at an excitation wavelength of 530 nm and an emission wavelength of 560 nm.

### Immunoblotting

Immunoblotting was performed as described (De et al. 2014). In brief, protein extracts from the cells were separated by 8 or 10% SDS-PAGE, transferred to PVDF membrane, blocked by 5% nonfat milk in TBS Tween 20 buffer (Thermo Fisher Scientific, cat. # 28360) and sequentially incubated with the indicated primary antibodies and horseradish peroxidase (HRP)-linked secondary antibody. SuperSignal West Pico PLUS Chemiluminescent Substrate (Thermo Fisher Scientific, cat. # 34577) was used for detection. Microsoft PowerPoint and Snip & Sketch were used to crop images from unprocessed images.

### Real-Time Quantitative PCR (RT-qPCR)

RT-qPCR was performed as described previously (De et al. 2014). cDNA was synthesized from total RNA, using random hexamers and SuperScript III. Real-time PCR was performed with PR1MA qMAX Green qPCR Mix (Midwest Scientific, cat. # PR2000-N-5000) in a CFX Opus 384 Real-Time PCR System (Bio-Rad). For genomic DNA (gDNA), gDNA was isolated using a DNeasy Blood & Tissue Kit (Qiagen, cat. # 69504) and qPCR was performed using gene-specific primers.

### RNA Immunoprecipitation (RIP)

RIP was performed using the Magna RIP™ RNA-Binding Protein Immunoprecipitation Kit (EMD Millipore, cat. # 17-700) according to the manufacturer’s instructions. Briefly, cells expressing FLAG-tagged WT METTL16 were lysed under conditions preserving RNA-protein interactions. An aliquot of lysate was reserved as input control. The remaining lysate was incubated with anti-FLAG antibody-conjugated magnetic beads to immuno-precipitate METTL16-RNA complexes. After washing, RNA was purified from immuno-precipitates and subjected to RT-qPCR analysis. Enrichment of target transcripts was calculated using the percent input (% input) method. Normal mouse IgG served as a negative control.

### m^6^A RNA Immuno-precipitation (MeRIP)-qPCR

This experiment was performed using the Magna MeRIP™ m^6^A Kit (EMD Millipore, cat. # 17-10499) according to the manufacturer’s instructions. Briefly, mRNA was purified from total RNA using the GenElute mRNA Miniprep Kit (Millipore Sigma, cat. # MRN10). Following treatment with DNase (DNA-*free* DNA Removal Kit, Invitrogen, cat. # AM1906), 3 μg of isolated mRNA was fragmented to approximately 100-200 nucleotides. A portion of fragmented mRNA was reserved as input control, and the remaining mRNA (∼2 μg) was incubated with anti-m^6^A antibody-conjugated magnetic beads to immuno-precipitate m^6^A-modified transcripts. After washing, methylated mRNA was eluted, purified, and reverse transcribed. Enrichment of specific transcripts was quantified by qPCR and calculated using the percent input (% input) method. Normal mouse IgG was used as a negative control. Primers for known m^6^A-modified (positive) and non-modified (negative) regions of EEF1A1 were included with the kit as controls for specificity of m^6^A immunoprecipitation.

### Taxane accumulation assay

Intracellular taxane accumulation was assessed using fluorescein-labeled paclitaxel (Flutax-2; Oregon Green™ 488 Taxol; Invitrogen, cat. # P22310). Cells (6 × 10^5^ per well) were seeded in 6-well plates and cultured overnight in RPMI 1640 medium supplemented with 5% FBS. Following PBS washing, cells were incubated with 500 nM Flutax-2 in Opti-MEM I reduced-serum medium at 37°C for 3 h. After incubation, the medium was removed, and cells were rinsed with 1 mL ice-cold PBS. Cells were then trypsinized and centrifuged at 2,500 rpm for 5 min. The cell pellets were washed three times with 1 mL ice-cold PBS to remove extracellular Flutax-2 and subsequently resuspended in 500 μL ice-cold PBS. Fluorescence intensity was measured using a BD FACSymphony A5 SE (5-laser configuration; BD Biosciences, San Jose, CA, USA). Flutax-2 was detected using a 537/32 nm bandpass (B537) filter (detection range 521-553 nm). At least 10,000 events were acquired per sample. Data were analyzed using FlowJo version 10.9. (BD Biosciences).

### Immunofluorescence Microscopy

Cells were cultured on 4 or 8-well chamber slides (EMD Millipore). After experimentation, cells were fixed in 4% paraformaldehyde (PFA) in PBS for 30 min, rinsed twice with PBS, and permeabilized with 0.1% Triton X-100 in PBS (PBST) for 15 min. Cells were then blocked with 10% goat serum in PBS for at least 1 h. For immunostaining, ABCB1 primary antibody (1:250) was applied for 2 h at room temperature, followed by three washes with PBST. Secondary antibodies conjugated to Alexa Fluor 488 or Alexa Fluor 555 (1:500) were then added. Slides were mounted with ProLong Gold Antifade containing DAPI (Invitrogen, cat. # P36941) and imaged using an EVOS M5000 microscope (Invitrogen).

### *In vitro* methyltransferase activity assay

Methyltransferase activity was assessed using the MTase-Glo^TM^ Methyltransferase Assay (Promega, Cat. No. V7601), which quantifies the formation of the post-methylation product S-adenosylhomocysteine (SAH) via luminesence. The MAT2A hp4, containing the consensus 5-UAC**A\***GAG methylation motif (Satterwhite and Mansfield 2022; Pendleton et al. 2017), was ordered as a synthetic RNA oligonucleotide (Integrated DNA Technologies, Coralville, IA, USA). Substrate RNA was reconstituted in nuclease-free water, heated at 65°C for 10 min, and then cooled to room temperature to allow proper annealing of the RNA secondary structure prior to use. METTL16^WT^ and METTL16^N184A^ were prepared as a serial dilution in Assay Buffer (10 mM TRIS pH = 7.9, 10 mM MgCl_2_, 150 mM KCl, 200 µM SAM) at 2X final concentration, and 4 µL was added to a black non-binding 384-well plate (Corning). 4 µL of 2X substrate RNA (10 µM final concentration) was added and the plate was incubated for 30 minutes at 37°C. 2 µL of MTase-Glo^TM^ Reagent was added to each well and incubated for 30 min at 37°C; then 2 µL of MTase-Glo^TM^ Detection Solution was added to each well and incubated for 30 min at 37°C, as suggested by the manufacturer. Luminescence was measured using a Cytation 5 microplate reader (BioTeK) and background luminescence from protein-free control wells was subtracted. All reactions were performed in technical triplicate and reported as mean ± SD from a representative of three biological replicates.

### Polysome fractionation followed by qPCR

Polysome profiling was performed to assess mRNA translational status. Cells were grown in two 15 cm dishes to 70–80% confluency and treated with cycloheximide (100 μg/mL) for 10-15 min at 37°C. Cells were then washed twice with ice-cold PBS containing 100 μg/mL cycloheximide, scraped into PBS supplemented with cycloheximide, and centrifuged at 3,500 rpm for 10 min at 4°C. Cell pellets were resuspended in 1 mL polysome lysis buffer [10 mM HEPES-KOH, pH 7.9, 100 mM KCl, 2.5 mM MgCl₂, 1 mM DTT, 0.1% NP-40, 100 U/mL RNase inhibitor, Roche cOmplete™ Protease Inhibitor Cocktail EDTA-free (Sigma Aldrich, cat. # 11873580001), and 100 μg/mL cycloheximide] and homogenized by passing through a 25-gauge needle on ice. Lysates were cleared by centrifugation at 6,000 rpm for 15 min at 4°C, and the concentration was determined by measuring absorbance at 260 nm. Equal amounts of clarified lysate were loaded onto linear 10-50% sucrose gradients prepared in polysome buffer (10 mM HEPES, pH 7.4, 100 mM KCl, 2.5 mM MgCl₂, and 1 mM DTT). Gradients were centrifuged at 4°C using a SW32.1 rotor (Beckman Coulter) at 17,000 rpm for 20 h. Following centrifugation, gradients were fractionated using ISCO Programmable Density Gradient System with continuous monitoring at 254 nm using an ISCO UA-6 absorbance detector, and fractions were collected for downstream RNA analysis. RNA was isolated from individual sucrose gradient fractions using Trizol reagent (Invitrogen, cat. # 15596018). Equal amounts of RNA from each fraction were reverse-transcribed, and qPCR was performed using gene-specific primers. Data represent the mean ± SD of triplicate samples.

### Vivo-Morpholino Treatment and Xenograft Studies

A translation-blocking human METTL16 -specific Vivo-Morpholino and standard control Morpholino were synthesized by GeneTools LLC (Philomath, OR). For *in vitro* Vivo-Morpholino treatment, MDA-MB-231-PacR cells were treated with METTL16-targeting Vivo-Morpholino or control Vivo-Morpholino at the indicated concentration. Vivo-Morpholinos were added directly to complete culture medium without the use of transfection reagents and incubated for the indicated times at 37°C. Cell survival was determined by lysing the cells with 1 mol/L NaOH and measuring absorbance at 260 nm. For *in vivo* xenograft studies, MDA-MB-231-PacR cells (1 × 10⁵ cells) were orthotopically implanted into the mammary fat pads of female NOD/scid/IL-2Rγ (NSG) mice. Once tumors reached approximately 10 mm³, mice were randomized into two groups and treated with either control Vivo-Morpholino or METTL16 Vivo-Morpholino at a dose of 200 µg per tumor per mouse. Treatments were administered for three doses. Tumor volumes were measured using digital calipers, and tumor growth ratios were calculated by normalizing tumor volume at each time point to baseline volume at treatment initiation. All animal experiments were conducted in accordance with institutional guidelines and approved protocols.

### Statistical analysis

Values were expressed as means ± SD. *P* values were based on the paired *t* test and the significance was set at 0.05 (marked with an asterisk wherever data are statistically significant).

## Results

### METTL16 promotes taxane resistance in TNBC

To identify novel drivers of taxane resistance in TNBC, we employed an unbiased insertional mutagenesis approach designed to identify proteins whose overexpression confers a survival advantage under taxane selection pressure. Using validation-based insertional mutagenesis (VBIM), which involves the use of lentiviral vectors, we randomly inserted the strong CMV promoter into the genome of the taxane-sensitive TNBC cell line MDA-MB-468 (De et al. 2009).

The cells were treated with 4 nM docetaxel and, after 10 days, a docetaxel-resistant clone was isolated. To further identify the mRNA target associated with resistance, we performed RNA-based cloning utilizing vector sequences within the bicistronic mRNA expressed from the inserted promoter (De et al. 2014), finding this mRNA encodes the RNA methyltransferase, METTL16. Full-length METTL16 has 562 amino acids (Ruszkowska 2021), but the truncated version lacks 42 residues at the N-terminus. Fortunately, the truncated METTL16 protein retains its catalytic domain, represented by a characteristic Rossmann fold, spanning residues 79 to 288, that is essential for binding to S-adenosylmethionine (SAM) and RNA (Ruszkowska et al. 2018). To confirm that METTL16 is a mediator of docetaxel resistance, full-length METTL16 cDNA was introduced into un-mutagenized TNBC cells (MDA-MB-468, MDA-MB-231, and HCC70), resulting in stable pools in which METTL16 is overexpressed (Fig. 1 A,B,C). These cells were significantly more resistant to docetaxel than the vector controls (Fig. 1 D,E,F). To determine whether high expression of METTL16 caused resistance to another taxane, we treated MDA-MB-468 and MDA-MB-231 cells overexpressing METTL16 with paclitaxel, finding that enhanced levels of this protein increased resistance similarly (Fig. 1 G,H). These results confirm the role of METTL16 in taxane resistance in TNBC cells, prompting us to investigate whether METTL16 is similarly upregulated and functionally required in TNBC models of acquired paclitaxel resistance.

**Fig 1.**
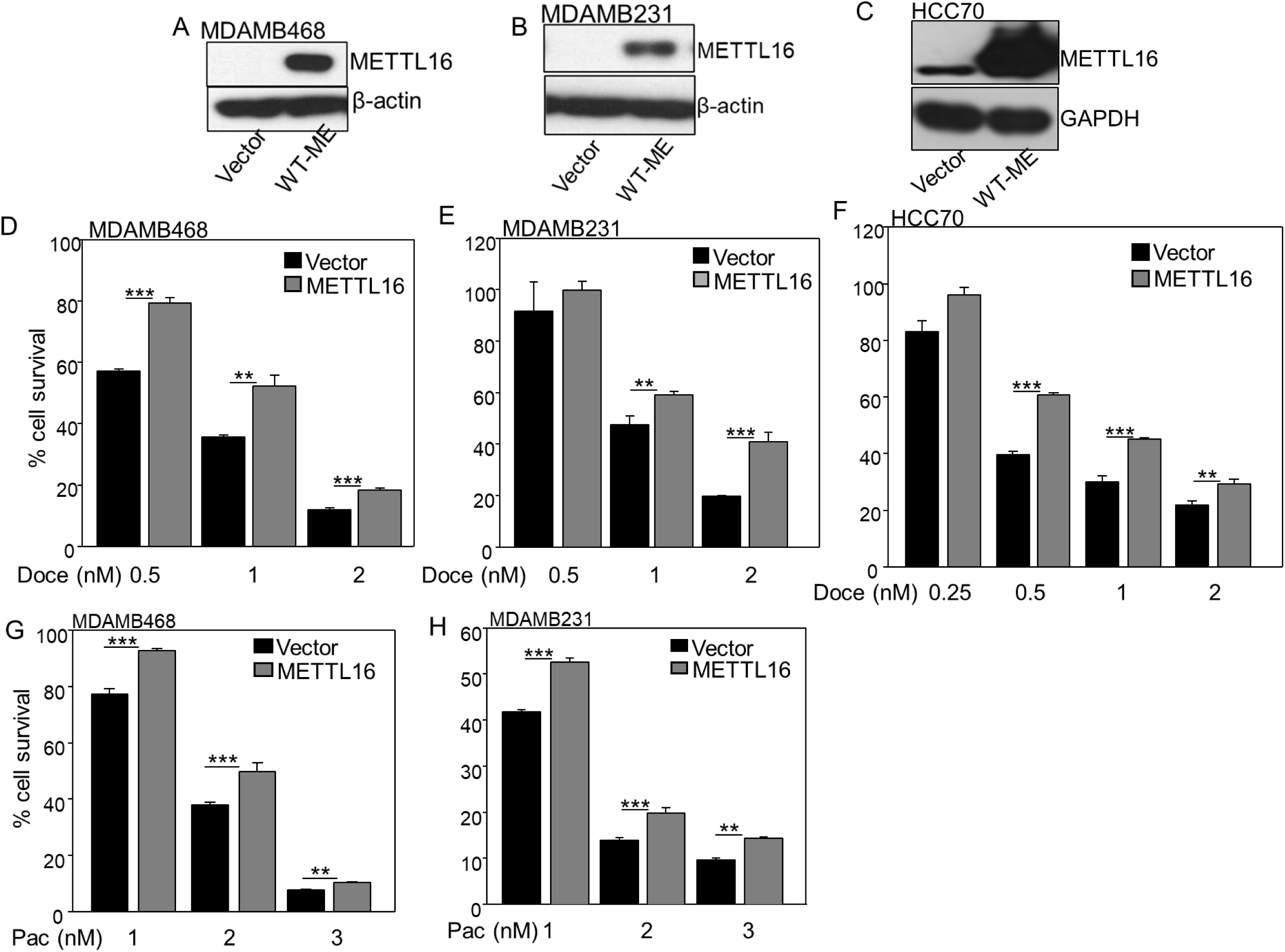
METTL16 overexpression mediates taxane resistance in TNBC cells. A, B, **C.** MDAMB468, MDAMB231 and HCC70 cells were infected with a vector encoding full-length METTL16 or empty vector and expression was examined by the Western method. **D, E, F, G, H.** Cell survival assays were performed following treatment with docetaxel (Doce; D, E, F) or paclitaxel (Pac; G, H) for 72h. The cells were lysed with 1 M NaOH and diluted 50-fold before the A260 was measured, as an indication of the total amount of nucleic acid. The fraction of surviving cells was determined by normalizing the data from taxane-treated cells to the data from DMSO controls. Results are represented by means ± SD. Data were analyzed using Student’s t-test. P values of <0.05 are considered statistically significant. **, P< 0.01; ***, P< 0.001.

### Down-regulation of METTL16 increases drug sensitivity in paclitaxel-resistant (PacR) TNBC cells with high endogenous METTL16

To determine whether METTL16 contributes to acquired paclitaxel resistance, we utilized paclitaxel-resistant (PacR) TNBC cell lines and examined METTL16 expression and function. The resistant cell lines MDA-MB-231-PacR and SUM159PT-PacR, generated by gradual paclitaxel exposure of the parental MDA-MB-231 and SUM159PT cells (Roberts et al. 2020), were maintained in 100 nM paclitaxel as a maintenance dose. METTL16 mRNA (Fig. 2 A,B) and protein (Fig. 2 C,D) levels were increased in the PacR cell lines compared to the paclitaxel-sensitive cells. Since gene amplification is a common mechanism by which cancer cells increase the expression of genes that promote survival or resistance to therapy (Matsui et al. 2013), we examined whether the increased expression of METTL16 in PacR TNBC cells is driven by changes in gene copy number. Quantitative RT-PCR (qPCR) analysis showed comparable gDNA abundance in the sensitive and PacR cells (Suppl. Fig. 1 A,B), indicating that increased METTL16 expression is driven by a transcriptional or post-transcriptional mechanism rather than gene amplification in these resistant cells. In addition to METTL16, other well-studied m^6^A RNA methyltransferases, including METTL3 and METTL14, play critical roles in promoting breast cancer (Wang et al. 2020; Yi et al. 2020). However, we did not observe any change in the expression levels of METTL3 and METTL14 in our PacR cells (Fig. 2 E,F,G,H).

**Fig 2.**
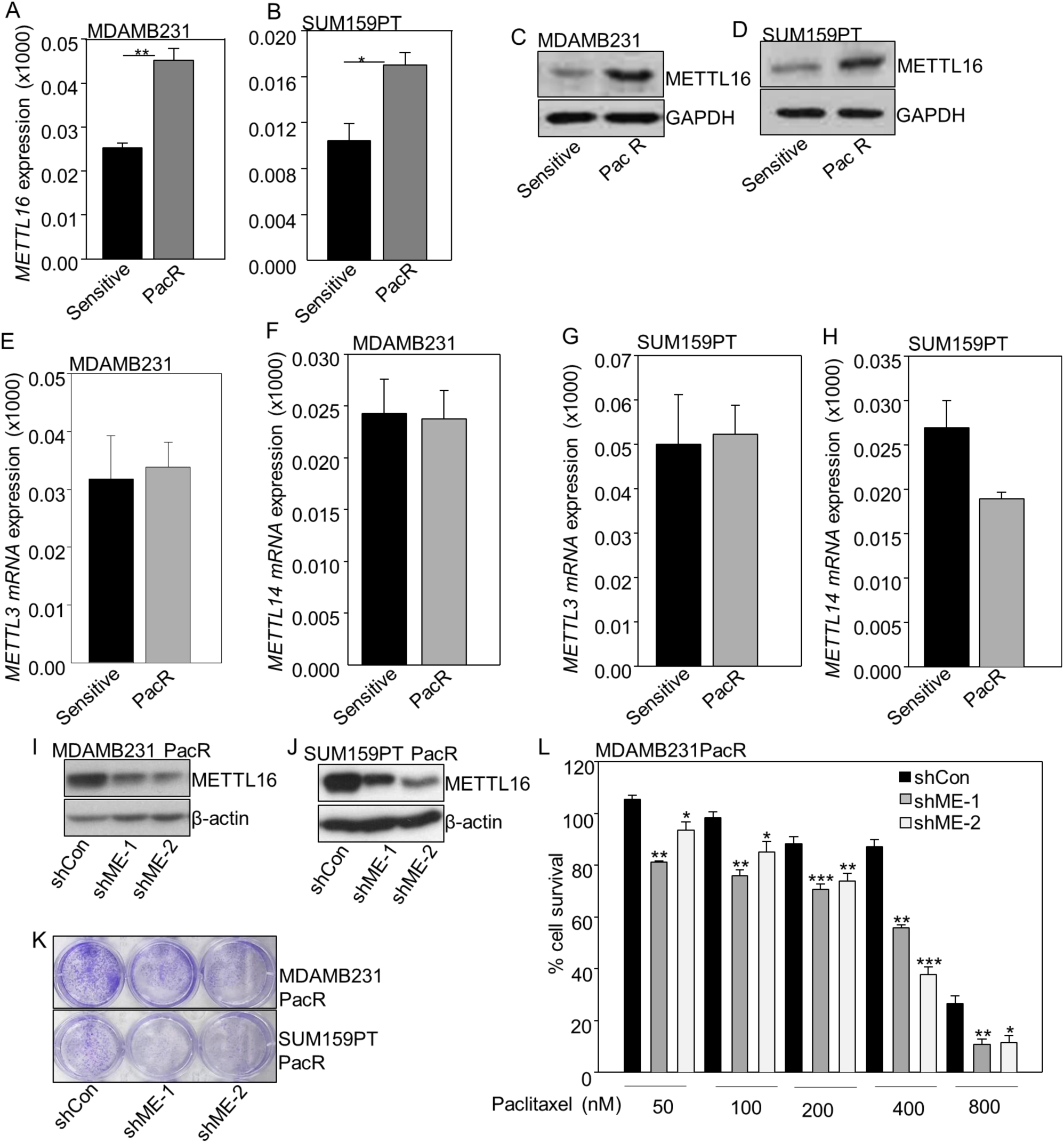
Down-regulation of METTL16 increases drug sensitivity in paclitaxel-resistant (PacR) TNBC cells. A, **B.** METTL16 mRNA expression level was determined by qPCR with 18 S RNA as a control. MDAMB231 PacR and SUM159PT PacR cells were generated from parental (sensitive) cells and are resistant to 100 nM paclitaxel. **C, D.** METTL16 protein levels were determined by Western analysis, using total cell lysates. GAPDH was used as a control. **E, F, G, H**. METTL3 and METTL14 mRNA expression in parental (sensitive) and PacR cells. **I, J, K, L**. Down-regulation of METTL16 decreases cell survival in untreated taxane-resistant TNBC cells. **I, J.** METTL16 expression was assayed by Western analysis in MDAMB231 PacR and SUM159PT PacR cells after knockdown of METTL16 (shME-1 and -2). Expression of non-targeting shRNA served as the control (shCon). **K.** Cell viability of the METTL16 knockdown and control cells was determined by staining with crystal violet. **L.** Cell survival assays were performed in MDAMB231 PacR cells with METTL16 knockdown and in control cells after treatment with paclitaxel. The cells were plated at 5 x 10^3^ cells/well in 96-well plates in triplicate. After 24 h, the cells were treated with the indicated concentrations of paclitaxel. Cell viability was determined after 5 days by CyQuant Direct assay and normalized to controls. Data are expressed as means ± SD. ***, P< 0.001; **, P< 0.01; *, P< 0.05.

To determine whether METTL16 is essential for paclitaxel resistance in our PacR cells, we down-regulated its expression using a METTL16-targeting shRNA (Fig. 2 I,J), observing a substantial decrease in cell survival, even without taxane treatment (Fig. 2 K). To determine whether METTL16 down-regulation could re-sensitize surviving PacR cells to paclitaxel, MDA-MB-231-PacR cells expressing METTL16-targeting shRNA were treated with increasing concentrations of paclitaxel (50-800 nM). We observed a marked increase in paclitaxel sensitivity in these cells, with a significant reduction in cell survival at concentrations as low as 50 nM, compared to control cells expressing a non-targeting shRNA (Fig. 2 L). Furthermore, we observed that PacR TNBC cells with high endogenous WT METTL16 expression levels have reduced sensitivity to the anthracycline doxorubicin (Suppl. Fig. 2), suggesting that METTL16 contributes to resistance to drugs other than taxanes known to be transported by ABCB1 (Piska et al. 2023). Together, these findings suggest that therapeutic targeting of METTL16 may be effective in overcoming paclitaxel resistance in patients with TNBC.

### The catalytic activity of METTL16 is required for taxane resistance

We assessed the impact of disrupting the catalytic activity of METTL16 by using a well-characterized inactive mutant. Residues within the ^184^NPPF^187^ sequence of METTL16 are essential for m^6^A methylation (Doxtader et al. 2018). We compared purified WT METTL16 and a protein with the inactivating mutation N184A by using the hairpin structure of MAT2A mRNA as a substrate, since METTL16-mediated m^6^A modification of MAT2A mRNA promotes its stability and expression (Zhang R 2022). WT METTL16 methylated the MAT2A substrate efficiently, consistent with prior reports (Flaherty et al. 2025). In contrast, the N184A mutant was much less active in this assay (Suppl. Fig. 3; Fig. 3 A), indicating that the N184A mutation inactivates METTL16. To determine whether the catalytic activity of METTL16 is required for taxane resistance, we found that MDA-MB-231 cells stably expressing the N184A mutant were significantly more sensitive to paclitaxel compared to control cells expressing an empty vector (Fig. 3 B). Consistently, in MDA-MB-231-PacR cells, stable overexpression of WT METTL16 further enhanced paclitaxel resistance, whereas expression of the N184A mutant failed to do so (Suppl. Fig. 4 A,B). Notably, N184A expression partially suppressed paclitaxel resistance in PacR cells but did not fully restore sensitivity, probably because of residual endogenous WT METTL16.

**Fig 3.**
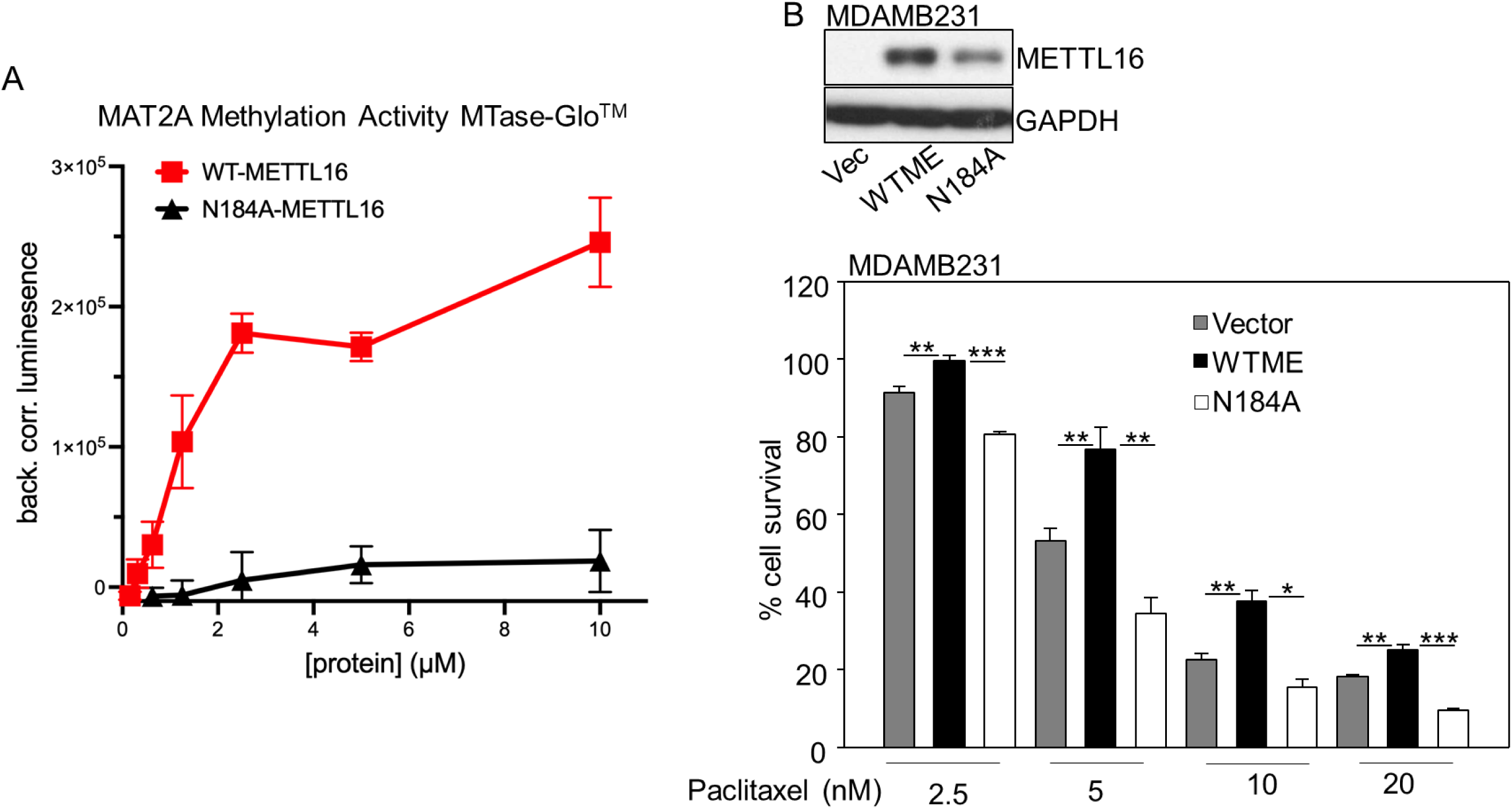
A point mutation in the catalytic domain of METTL16 increases the sensitivity to taxanes in TNBC cells. **A.** Recombinant protein biochemical methylation assay (MTase-Glo^TM^) showing that recombinant WT METTL16 exhibits robust, dose-dependent m^6^A methylation activity toward the MAT2A substrate, whereas the catalytically inactive N184A mutant displays markedly reduced activity, as measured by background-corrected luminescence. **B.** MDAMB231 cells were infected with empty vector or lentiviral constructs encoding WT METTL16 (WT ME) or the N184A point mutation of METTL16. Upper panel: The level of METTL16 protein was determined by Western analysis of total cell lysates. GAPDH was used as a control. Lower panel: Cell survival assays were performed after treatment with paclitaxel at different doses. The fraction of surviving cells was determined by normalizing the data from paclitaxel treated cells to DMSO controls. **, P< 0.01; ***, P< 0.001.

### METTL16 upregulates ABCB1 (P-glycoprotein) expression and reduces intracellular paclitaxel accumulation

ABCB1, also known as P-glycoprotein, is a membrane transporter that plays a central role in multidrug resistance by actively exporting chemotherapeutic agents, including taxanes (Tian et al. 2023). Elevated ABCB1 expression in tumors is strongly associated with paclitaxel resistance, treatment failure, and disease relapse (Alalawy 2024). Consistent with this role, we observed significantly increased ABCB1 mRNA and protein levels in paclitaxel-resistant MDA-MB-231-PacR and SUM159PT-PacR cells compared with their paclitaxel-sensitive parental counterparts (Fig. 4 A,B). Although gene amplification has been reported as a conserved mechanism driving ABCB1 upregulation in paclitaxel-resistant pancreatic cancer (Bergonzini et al. 2024), quantitative genomic DNA analysis did not reveal altered/increased ABCB1 copy number in our PacR MDA-MB-231 and SUM159PT cell lines (Suppl. Fig. 5 A,B), indicating that ABCB1 upregulation in these cells occurs through post-genomic regulatory mechanisms rather than gene amplification.

**Fig 4.**
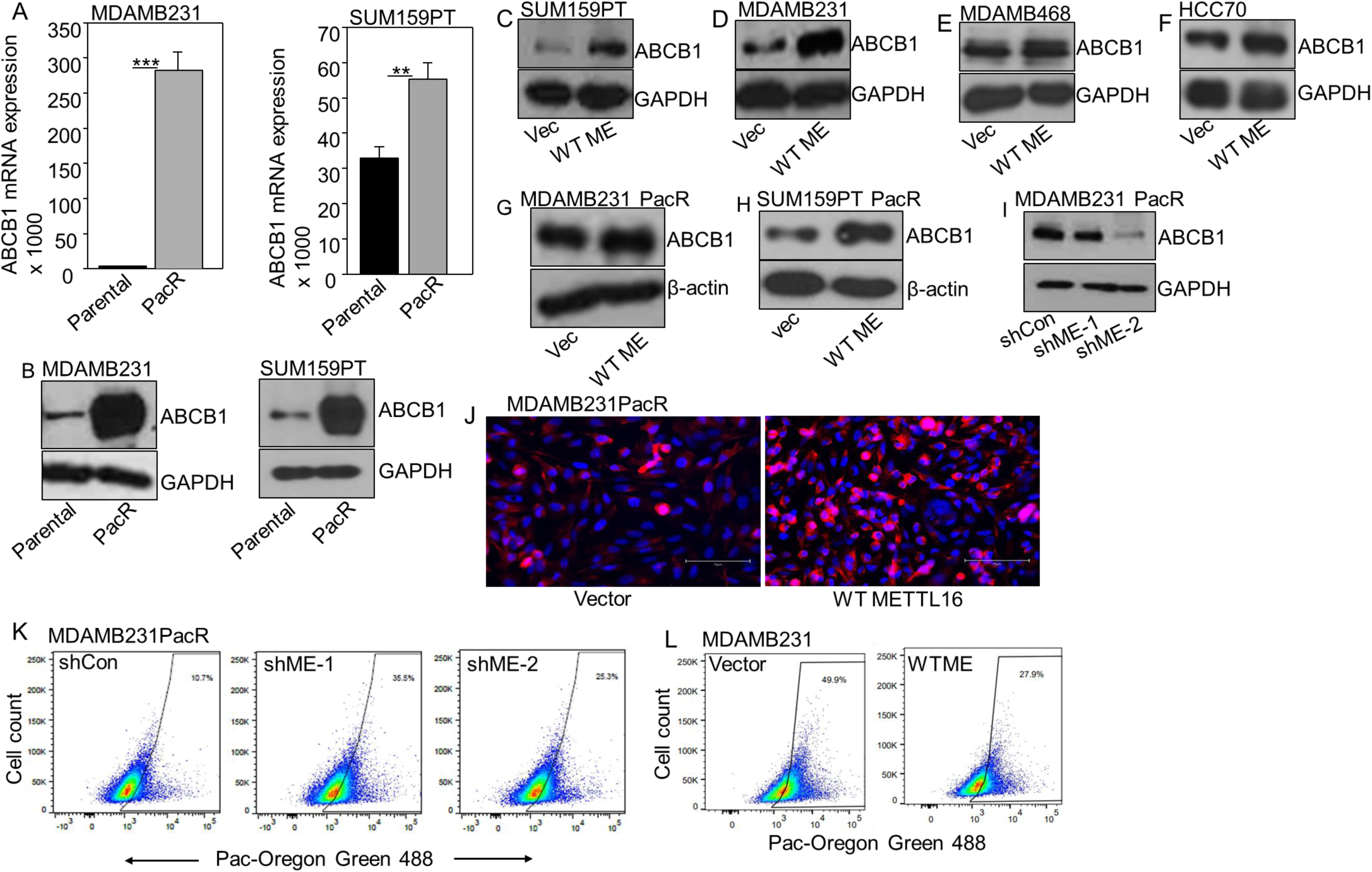
METTL16 upregulates ABCB1 (P-glycoprotein) and reduces intracellular paclitaxel accumulation. **A.** The mRNA expression of ABCB1 was determined by qPCR in parental (sensitive) and PacR MDAMB231 and SUM159PT cells. **B.** ABCB1 protein levels were determined by Western analysis in these cells, with GAPDH as a loading control. **C, D, E, F**. ABCB1 protein levels were detected in TNBC cells (SUM159PT, MDAMB231, MDAMB468, HCC70) overexpressing METTL16 (WT ME) or in control cells with empty vector (Vec). GAPDH was used as a loading control. **G, H**. ABCB1 protein levels were determined in the MDAMB231 PacR and SUM159PT PacR cells overexpressing WT ME or Vec. β-actin was used as a loading control. **I.** ABCB1 protein levels were determined in the MDAMB231 PacR cells with METTL16 knockdown (shME-1 and shME-2). GAPDH was used as a loading control. **J.** Immunofluorescence staining of ABCB1 in TNBC cells. Representative images of MDAMB231 PacR cells with vector control (left) and overexpressing WT METTL16 (right) stained for ABCB1 (red) and nuclei (DAPI, blue). Scale bar: 75 µm. **K, L.** Paclitaxel accumulation assay using flow cytometry. Paclitaxel-Oregon green fluorescence was measured to assess intracellular drug accumulation in (**K**) MDAMB231 PacR cells expressing control shRNA (shCon) or METTL16-targeting shRNAs (shME-1 and shME-2), and (**L**) in taxane-sensitive MDAMB231 cells with empty vector (Vector) or overexpressing WT METTL16 (WTMETTL16) following treatment with Flutax-2 (500 nM) for 3 h. Increased fluorescence indicates increased intracellular drug accumulation.

To determine whether METTL16 regulates ABCB1 expression, we overexpressed WT METTL16 in several TNBC cell lines, including SUM159PT, MDA-MB-231, MDA-MB-468, and HCC70 cells, observing a marked increase in ABCB1 protein levels (Fig. 4 C-F). Similarly, ABCB1 protein expression was further elevated in both MDA-MB-231-PacR and SUM159PT-PacR cells overexpressing WT METTL16 (Fig. 4 G,H). Notably, this increase in ABCB1 protein was not accompanied by a corresponding increase in ABCB1 mRNA levels in either paclitaxel-sensitive or PacR cells (Suppl. Fig. 5 C,D), suggesting that METTL16 regulates ABCB1 primarily at a post-transcriptional level. Conversely, shRNA-mediated down-regulation of METTL16 in MDA-MB-231-PacR cells led to a substantial reduction in ABCB1 protein expression (Fig. 4 I). To further validate the effect of WT METTL16 on ABCB1 expression, immunofluorescence analysis of MDA-MB-231-PacR cells confirmed increased ABCB1 protein levels in WT METTL16 overexpressing cells compared with vector controls (Fig. 4J). To assess whether METTL16-mediated regulation of ABCB1 extends beyond TNBC, we down-regulated METTL16 in A549 lung adenocarcinoma cells, which also express ABCB1. Consistent with our TNBC findings, METTL16 knockdown resulted in a significant decrease in ABCB1 protein levels (Suppl. Fig. 5 E), indicating that METTL16-dependent regulation of ABCB1 is conserved across different cancer types.

Using Flutax-2 (fluorescent paclitaxel) accumulation assays, we found that down-regulation of METTL16 in MDA-MB-231-PacR cells increased Flutax retention (∼35% or 25% Flutax-positive cells) compared to non-targeting controls (∼10% Flutax-positive cells), indicating a substantially reduced drug efflux capacity (Fig. 4K). Conversely, overexpression of WT METTL16 in paclitaxel-sensitive MDA-MB-231 cells decreased Flutax accumulation (∼28% Flutax-positive cells) relative to empty vector controls (∼50% Flutax-positive cells) (Fig. 4L). These findings indicate that METTL16 promotes taxane resistance by upregulating ABCB1 expression and enhancing drug efflux, thereby reducing efficiently intracellular paclitaxel accumulation and limiting its cytotoxic effects. To directly assess functional ABCB1 transport activity, we performed a Calcein-AM efflux assay using MDA-MB-231-PacR cells overexpressing WT METTL16, with or without the ABCB1 inhibitor verapamil (Frayde-Gómez et al. 2026). Verapamil treatment resulted in a rightward shift in fluorescence intensity and an almost two-fold increase in median fluorescence intensity (MFI: 17,403 vs. 9,086), indicating increased intracellular calcein retention (Suppl. Fig. 6). These results confirm active ABCB1-mediated efflux in WT METTL16 PacR cells and confirm the reduced Flutax accumulation observed in the resistant phenotype.

Because paclitaxel exerts its cytotoxic effects by inducing apoptotic cell death (Abu Samaan et al. 2019), we next examined whether METTL16 modulates paclitaxel-induced apoptosis. Annexin V/PI staining was performed following paclitaxel treatment of MDA-MB-231-PacR cells expressing control or METTL16-targeting shRNAs. In control cells, paclitaxel exposure resulted in only a small decrease in live cells, from 66.0 % to 64.6 %, consistent with the resistant phenotype (Suppl. Fig. 7 A). In contrast, down-regulation of METTL16 enhanced paclitaxel-induced apoptosis. Live cells decreased from 72.8-74.6% in untreated METTL16-depleted cells to 55.4% with shME-1 and 63.9% with shME-2 following paclitaxel treatment, accompanied by a corresponding decrease in viability (Suppl. Fig. 7 B,C). These findings indicate that depletion of METTL16 sensitizes PacR TNBC cells to taxane-induced cell death by promoting progression to apoptosis. Together, the Flutax accumulation, Calcein-AM efflux, and Annexin V/PI apoptosis assays demonstrate that METTL16 enhances ABCB1 transport activity, thereby reducing intracellular paclitaxel levels and apoptotic responses.

### METTL16 binds to ABCB1 mRNA, enhancing its m^6^A methylation

RNA immunoprecipitation assays (RNA-IP) in MDA-MB-231-PacR cells overexpressing FLAG-tagged WT METTL16 confirmed its robust expression (Fig. 5A, upper panel) and resulted in a substantial enrichment of ABCB1 mRNA, compared with controls (Fig. 5A, lower panel-left). Enrichment of MAT2A mRNA, a well-established direct substrate of METTL16, was also confirmed in FLAG-tagged WT METTL16 immuno-precipitates, validating the specificity and functional integrity of the assay (Fig. 5 A, lower panel-right). These data establish ABCB1 mRNA as an RNA target of METTL16 in taxane-resistant TNBC cells and is consistent with a post-transcriptional regulatory role for METTL16, in line with its function as an m^6^A RNA methyltransferase. Having established METTL16-ABCB1 interaction, we next investigated whether METTL16 regulates ABCB1 through m^6^A RNA methylation by performing m⁶A RNA immunoprecipitation followed by qPCR (MeRIP-qPCR). Putative METTL16-target methylation sites were identified by searching the ABCB1 mRNA sequence (NM_001348945.2) for the previously described METTL16-target consensus sequence, 5’-UAC<u>A</u>GAGAA-3’, where the underlined A is methylated (Pendleton et al. 2017; Yoshinaga et al. 2022). Since no identical sequences were found, we focused on sequences with no more than three mismatches and with a conserved A in the fourth position. To further narrow down and increase confidence in these sites, we cross-referenced the sequences with those identified in the online database RMBase v3.0 (https://rna.sysu.edu.cn/rmbase3/index.php) and the online Sequence-based RNA Adenosine Methylation Site Predictor (SRAMP, http://www.cuilab.cn/m6asiteapp/old). Both RMBase and SRAMP utilize the consensus sequence for the METTL3-METTL14 complex, which differs significantly from that of METTL16: 5’-DRACH-3’ or 5’-RRACH-3’, where D = A/G/U, R = A/G, and H = A/C/U (Xuan et al. 2018; Zhou et al. 2016). We reasoned that METTL16 consensus sequences located within or adjacent to the sequences identified by RMBase or SRAMP were likely to be located in RNA secondary structures that would be accessible to METTL16 as well as the METTL3-METTL14 complex. Following this analysis, we identified 3 adenine residues with the highest confidence to be modified: A2368, A4141, and A4497. The first two are within the coding sequence and the final one is in the 3’-UTR. In MAT2A mRNA, the METTL16-target consensus sequences are located within hairpin structures (Parker et al. 2011). To confirm our RMBase and SRAMP analyses, we entered the complete ABCB1 mRNA sequence into the RNAfold Web Server (http://rna.tbi.univie.ac.at/cgi-bin/RNAWebSuite/RNAfold.cgi) in order to predict whether or not our putative METTL16-target sequences were located within potential hairpin structures (Gruber et al. 2008). All three identified adenine residues were probably within or immediately adjacent to hairpin structures (Suppl. Fig. 8). Following this analysis, primers were designed to target these three regions of the METTL16 mRNA. By MeRIP-qPCR analysis, we demonstrated a significant increase in m^6^A enrichment on ABCB1 mRNA in WT METTL16-expressing MDA-MB-231-PacR cells compared with empty vector controls. Enhanced m^6^A modification was observed across all three target regions, indicating that METTL16 broadly promotes m^6^A deposition on ABCB1 RNA. IgG immunoprecipitation showed minimal signal, confirming the specificity of the m^6^A pull-down (Fig. 5 B-D). Furthermore, MeRIP-qPCR analysis showed a significantly reduced m^6^A enrichment on ABCB1 transcripts in MDA-MB-231-PacR cells with METTL16 down-regulation compared to the control cells (Fig. 5 E). These findings indicate that endogenous METTL16 is required for maintaining m^6^A modification of ABCB1 mRNA in PacR cells. To validate the efficiency and specificity of the MeRIP assay, we assessed m^6^A enrichment using positive and negative control primers targeting Eukaryotic Translation Elongation Factor 1 Alpha 1 (EEF1A1), a well-established, abundantly expressed housekeeping mRNA that contains m^6^A modifications, making it an ideal technical control for MeRIP assays (Magna MeRIP^TM^ m^6^A Kit, EMD Millipore). In MDA-MB-231-PacR cells expressing WT METTL16, MeRIP-qPCR revealed a significant increase in m^6^A enrichment on the EEF1A1 positive control region known to be m^6^A-methylated, whereas no enrichment was detected for the negative control region of EEF1A1 lacking m^6^A. IgG immunoprecipitation showed minimal signal for both positive and negative control RNAs (Suppl. Fig. 9). This result confirms successful and specific immunoprecipitation of m^6^A-modified RNA and demonstrates that WT METTL16 expression does not induce nonspecific background pull-down, validating the integrity of the MeRIP assay used to assess METTL16-dependent m^6^A modification of the target transcript, ABCB1. Together, the results demonstrate that METTL16 binds to and methylates ABCB1 mRNA, providing a basis for METTL16-mediated upregulation of ABCB1 expression through m^6^A modification.

**Fig 5.**
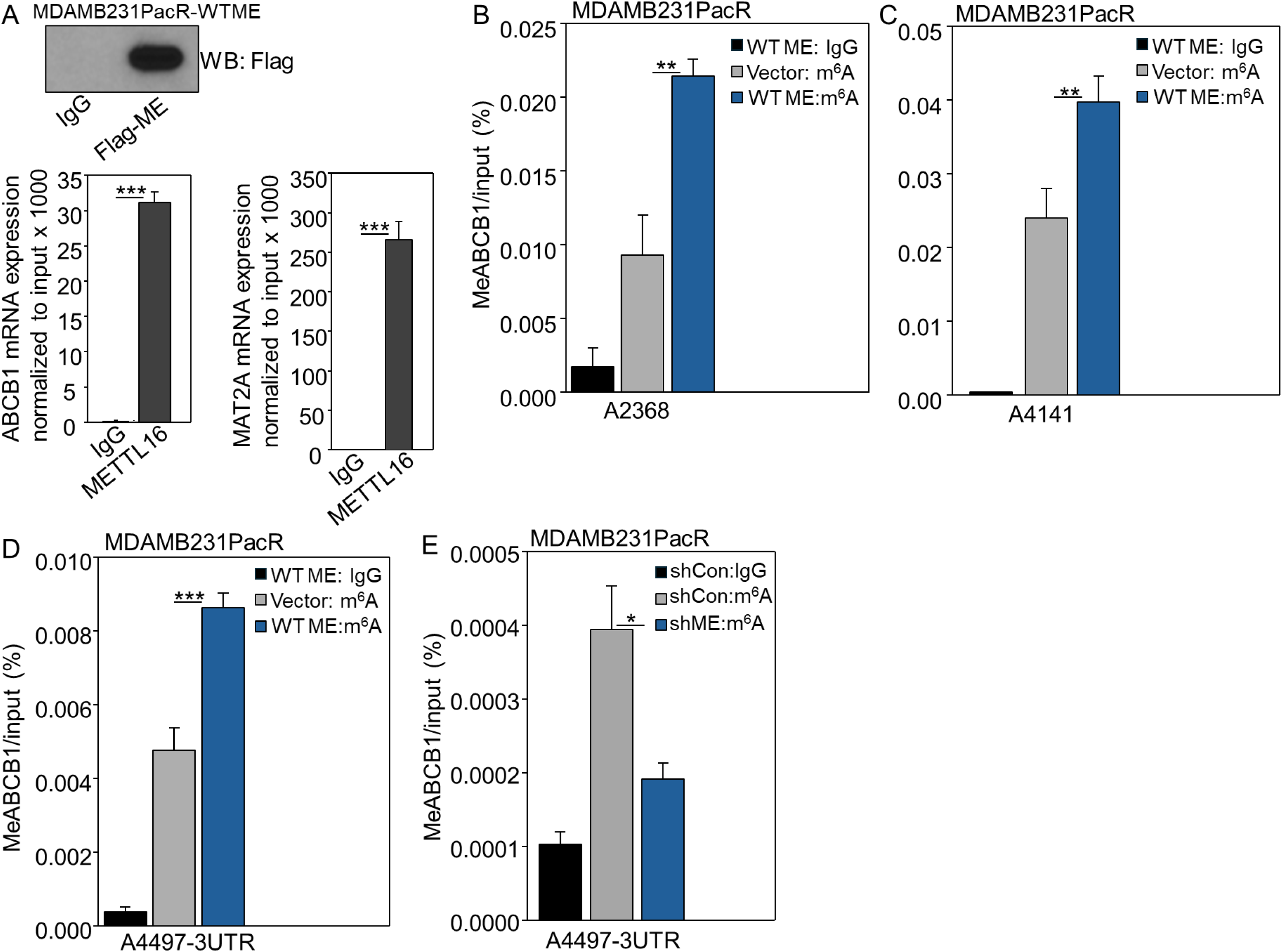
METTL16 binds to ABCB1 mRNA and enhances its m⁶A methylation in PacR TNBC cells. **A.** METTL16 RNA immunoprecipitation (RNA IP) reveals its association with ABCB1 mRNA. MDAMB231 PacR cells were transfected with FLAG-tagged WT METTL16 (Flag-ME). RNA IP was performed by using an anti-FLAG or control IgG antibody to pull down METTL16-bound RNAs, followed by qPCR analysis to detect ABCB1 or MAT2A mRNA. Upper panel: Immunoblot of the RNA IP fraction probed with anti-Flag antibody confirming efficient pull-down of Flag-METTL16. Lower panel: Enrichment of ABCB1 or MAT2A mRNA in the anti-FLAG immuno-precipitate, normalized to input. **B, C, D.** To determine m⁶A enrichment on ABCB1 mRNA, the Magna MeRIP m^6^A Kit (EMD Millipore) was used to perform m⁶A RNA immunoprecipitation followed by qPCR (MeRIP-qPCR) in MDAMB231 PacR cells overexpressing WT METTL16 (WT ME) or empty vector (Vector). Normal mouse IgG was used as a control. **E.** MeRIP-qPCR in MDAMB231 PacR cells with down-regulated METTL16 (shME) compared to non-targeting controls (shCon). Normal mouse IgG was used as a control.

### METTL16 methyltransferase activity promotes ABCB1 mRNA translation

Given the methylation of ABCB1 mRNA by METTL16, we investigated whether METTL16 enhances ABCB1 expression by promoting its translation. Polysome profiling was performed in MDA-MB-231-PacR and paclitaxel-sensitive cells expressing empty vector, WT METTL16, or the catalytically inactive N184A mutant. Cell lysates were separated on 10-50% sucrose gradients, fractionated, and abundance of ABCB1 mRNA in each fraction was analyzed by RT–qPCR. GAPDH or β-actin mRNAs were used as internal controls. Because the polysome:monosome (P:M) ratio provides a quantitative measure of translation efficiency, with higher P:M ratios reflecting increased ribosome loading and active protein synthesis, we compared P:M ratios across these conditions to assess whether METTL16 enhances ABCB1 translation. We calculated the P:M ratio by summing the qPCR (ΔCt) values for polysome fractions (fractions 9-14) and normalizing to the monosome fractions (fractions 6 or 6-7, wherever indicated).

In MDA-MB-231-PacR cells, ABCB1 mRNA in vector control was predominantly enriched in the monosome fraction with a P:M ratio of 2.5 (Fig. 6 A, left panel), indicating limited ribosome loading and reduced translational efficiency of ABCB1 mRNA under basal conditions. In contrast, expression of WT METTL16 resulted in a major shift of ABCB1 mRNA from monosomal to polysomal fractions with an increase of P:M ratio to 4.4, consistent with enhanced ribosome recruitment and active translation. Expression of the catalytically inactive N184A mutant reversed this effect. Under these conditions, ABCB1 mRNA showed substantially reduced polysomal association compared with WT METTL16 overexpressing cells (P:M ratio of 4.81 for mutant versus 6.41 for WT), and overall ABCB1 mRNA distribution between different fractions was more similar to that of the vector control (P:M ratio of 3.77 for vec; Fig. 6 B, left panel). This information indicates that METTL16 catalytic activity is required for maximal enhancement of ABCB1 translation, while loss of methyltransferase activity compromises ribosome loading. Importantly, GAPDH mRNA distribution was unchanged across all conditions, confirming that the observed shift was specific to ABCB1 and was not due to global translational alterations or fractionation artifacts (Fig. 6 A,B, right panels).

**Fig 6.**
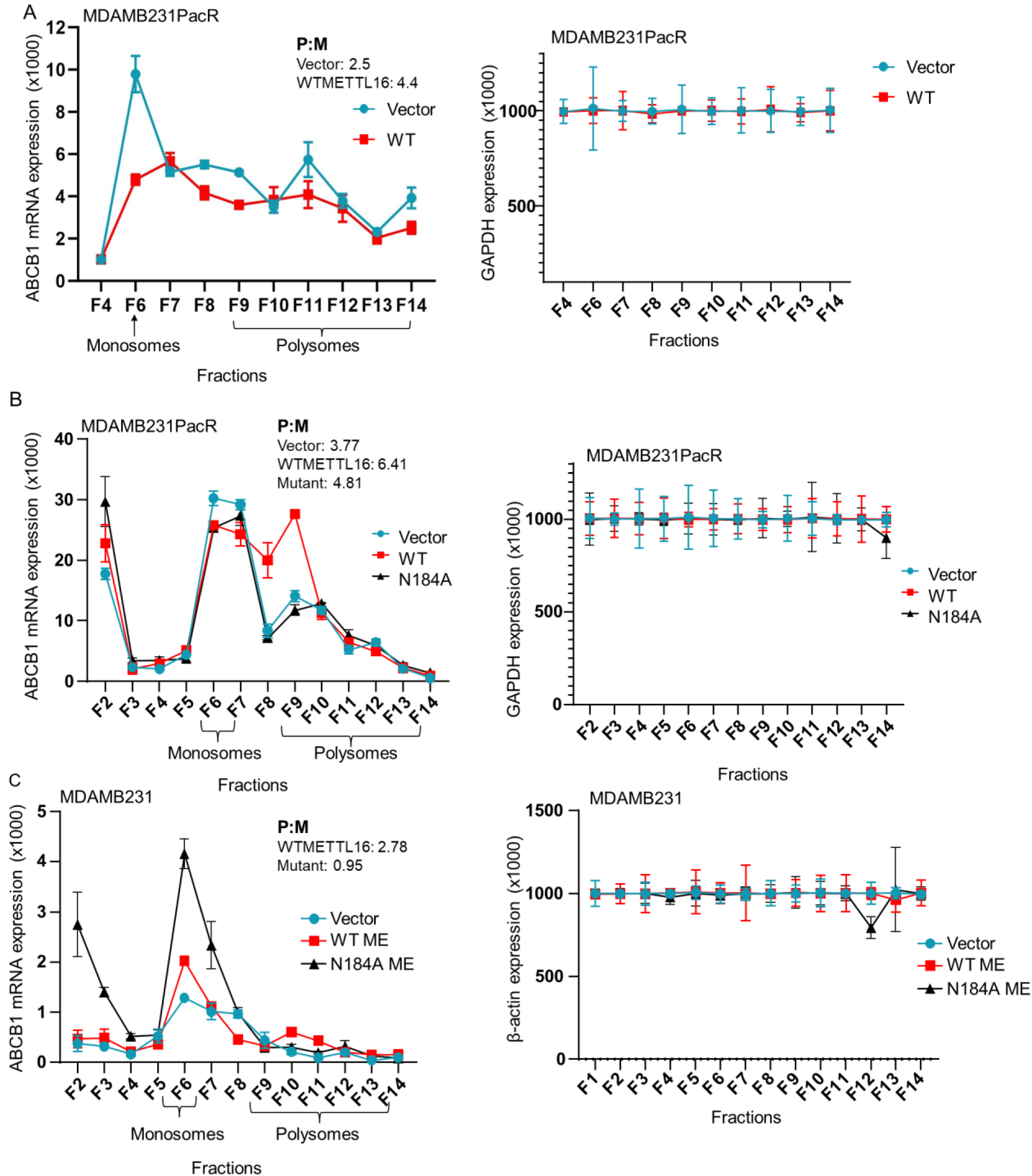
METTL16 enhances ABCB1 mRNA translation efficiency. A, B, **C**. Polysome profiling followed by qPCR was performed to examine the distribution of ABCB1 mRNA (left panels) and GAPDH or β-actin mRNA (right panels) across ribosomal fractions. Analyses were conducted in: (**A**) MDAMB231 PacR cells overexpressing WT METTL16 or an empty vector control; (**B**) MDAMB231 PacR cells overexpressing WT METTL16, the catalytically inactive METTL16 mutant (N184A), or an empty vector control; (**C**) taxane-sensitive MDAMB231 cells overexpressing WT METTL16, the N184A mutant, or an empty vector control. The P:M ratio was calculated by summing qPCR (ΔCt) values from polysome fractions (fractions 9-14) and normalizing to monosome fractions (fraction 6 for **A**, fractions 6 and 7 for **B, C**).

Similar analyses in MDA-MB-231 sensitive cells revealed that cells expressing the N184A mutant showed a substantial increase in monosomal ABCB1 mRNA compared to WT METTL16 overexpressing cells (Fig. 6 C, left panel). The P:M ratio was 2.78 for WT cells and 0.95 for the mutant cells, reflecting markedly impaired ribosome recruitment with the mutant. β-actin mRNA distribution was similar between the groups (Fig. 6 C, right panel). Together, the polysome distributions and P:M ratios in both PacR and paclitaxel-sensitive cells demonstrate that METTL16 enhances ABCB1 translation in a manner that is dependent on its methyltransferase activity, establishing a mechanistic link between METTL16-mediated m^6^A modification and ABCB1 translational regulation associated with taxane resistance.

### Clinical relevance of the METTL16-ABCB1 axis in TNBC

To examine whether METTL16 expression is clinically relevant in TNBC patients treated with chemotherapy, we performed a Kaplan-Meier survival analysis (Győrffy 2021) using publicly available patient datasets. In a cohort of systemically untreated TNBC patients (n = 43), stratified by median METTL16 expression (Affy ID: 226744_at), higher METTL16 levels were associated with reduced relapse-free survival (RFS); however, this association did not reach statistical significance (hazard ratio [HR] = 3.65, 95% confidence interval [CI]: 0.76-17.56; *log-rank P* = 0.084; Suppl. Fig. 10 A). We also analyzed a chemotherapy-treated TNBC cohort with explicitly annotated treatment status (“include all” setting; n = 50). METTL16 expression demonstrated significant prognostic value in this group. Patients with high METTL16 expression exhibited significantly shorter RFS compared to those with low expression (HR = 5.4, 95% CI: 1.54-19.02; *P* = 0.0032; Suppl. Fig. 10 B). These clinical associations support a role for METTL16 in therapy resistance *in vivo*.

To assess the clinical relevance of the METTL16-ABCB1 axis, we examined the correlation between METTL16 and ABCB1 expression in TNBC patient cohorts using Breast cancer-GenExMiner, an integrated bioinformatics platform that analyzes normalized gene expression data from publicly available breast cancer transcriptomic datasets to identify correlations between the expression of genes across TNBC molecular subtypes. Pearson’s correlation analysis across all TNBC samples revealed a modest but significant positive association between METTL16 and ABCB1 expression (r = 0.22, p = 0.0001; n = 293), indicating that higher METTL16 levels are associated with increased ABCB1 expression in tumors (Suppl. Fig. 11 A). We next evaluated the relationship between METTL16 and ABCB1 expression across different TNBC subtypes. Notably, ABCB1 mRNA expression was significantly higher in the mesenchymal-like immune-altered (MLIA) subtype compared with other TNBC subtypes, such as basal-like immune-activated (BLIA), basal-like immune-suppressed (BLIS), and luminal androgen receptor (LAR) (Jézéquel P, 2024; Suppl. Fig. 11 B). Importantly, a strong positive correlation between METTL16 and ABCB1 expression was observed specifically within the MLIA subtype (r = 0.48, p < 0.0001; n = 121) (Suppl. Fig. 11 C). This subtype is characterized by mesenchymal features, altered immune signaling, enrichment of stem-like properties, and marked resistance to chemotherapy, radiotherapy, and immunotherapy (Jézéquel et al. 2024). Consistent with this classification, MDA-MB-231 and SUM159PT are categorized as mesenchymal stem-like TNBC models (Huang et al. 2024), directly linking our experimental systems to this clinically aggressive and therapy-resistant patient subgroup. Given the well-established role of ABCB1 in mediating multidrug efflux, its elevated expression in MLIA tumors probably contributes to the pronounced therapeutic resistance observed in this subtype. Together, these clinical correlations provide a strong rationale for *in vivo* validation of METTL16 as a therapeutic target in TNBC.

### METTL16-specific Vivo Morpholino inhibits cell survival and tumor growth in PacR TNBC

To evaluate the therapeutic potential of METTL16 inhibition, we employed a Vivo-Morpholino, synthetic antisense oligonucleotides designed to block METTL16 translation initiation. Treatment of MDA-MB-231-PacR cells with the METTL16-targeting morpholino (5 µM, 48 h) resulted in marked down-regulation of METTL16 protein as assessed by Western blot (Fig. 7 A), accompanied by a significant decrease in cell viability (***P < 0.001), with over 50% reduction in survival compared to cells treated with control oligo (Fig. 7 B). To test *in vivo* efficacy, PacR TNBC xenografts in NSG mice were treated intratumorally with either METTL16 Vivo-Morpholino or control oligo. Tumor volumes were measured over time and normalized to baseline values at treatment initiation. As shown in Fig. 7 C, the morpholino-treated group exhibited a significant reduction in tumor growth over the 8-day period following treatment, with ∼2-fold lower tumor growth ratios after three doses. These results establish METTL16 as a therapeutically actionable vulnerability in PacR TNBC.

**Fig 7.**
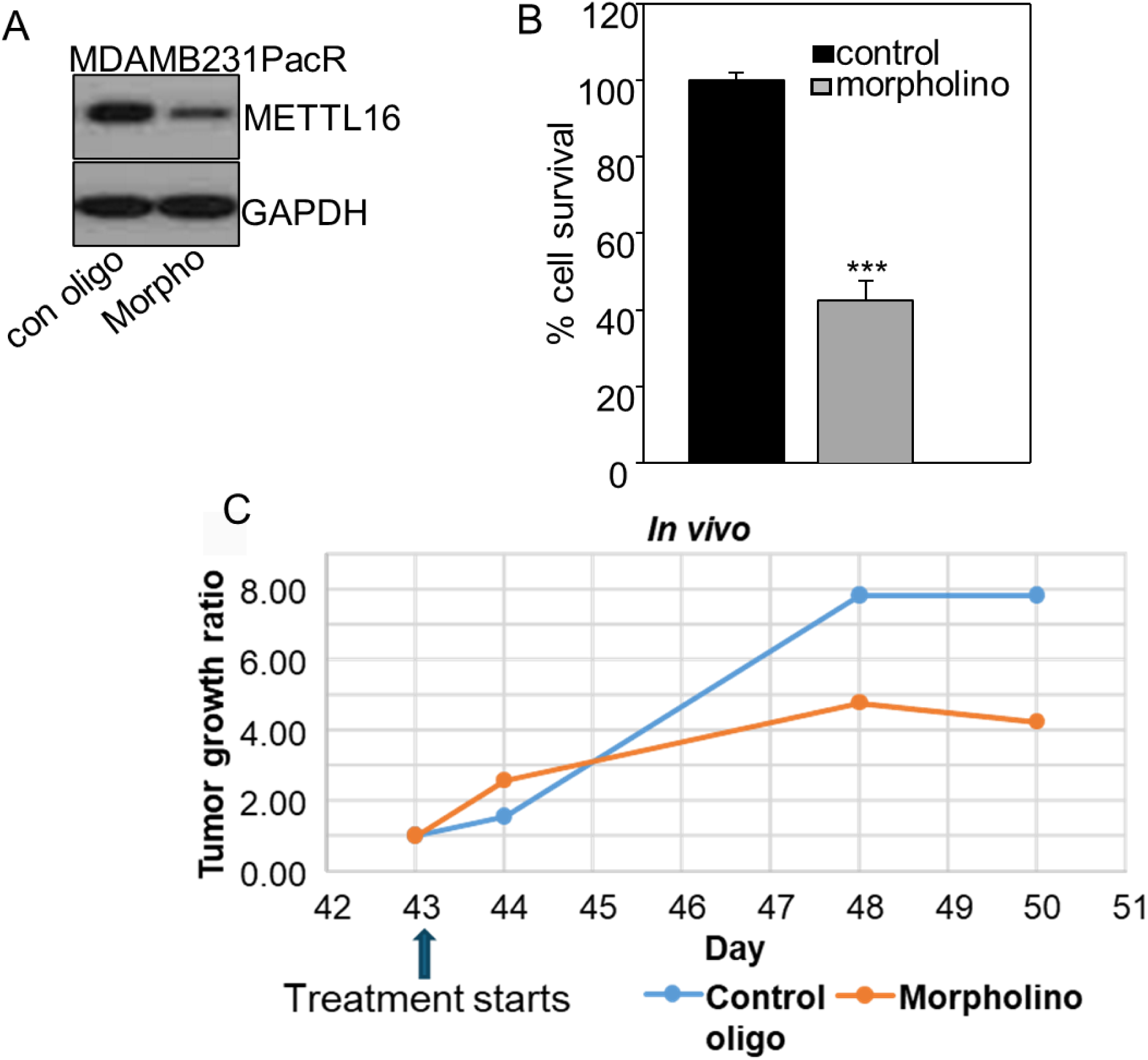
METTL16-targeting morpholinos inhibit tumor growth in PacR TNBC cells. **A.** METTL16 protein level determined by Western analysis in MDAMB231 PacR cells after treatment with Vivo-Morpholino (5 µM) for 48h. **B.** Cell survival in Vivo-Morpholino-treated cells determined by AlamarBlue assay. ***, P< 0.001. **C.** Tumor growth in mice was assessed by caliper measurement in control and morpholino-treated groups (n=5). The tumor growth ratio was normalized by setting the ratio to 1 for both groups on the day of treatment initiation. The growth ratio was calculated as the tumor volume at each time point divided by the baseline volume at treatment initiation. The data demonstrate inhibition of tumor growth in the morpholino-treated group compared to control after three treatments.

### METTL16 is critical for maintaining TNBC cell survival

A previous report demonstrated that complete genetic ablation of METTL16 using CRISPR–Cas9 induced cell death in hepatocellular carcinoma cells, whereas METTL16 knockout had no significant effect on the viability of normal hepatocytes (Su et al. 2022). Using the same CRISPR constructs, we similarly observed that METTL16 knockout was not tolerated in MDA-MB-231 cells, resulting in an approximately 80% reduction in cell survival compared with control cells (Fig. 8 A,B,C). Similar results were observed in an additional TNBC cell line, SUM159PT, in which METTL16 knockout led to a ∼70% decrease in cell survival relative to controls (Fig. 8 D,E). Consequently, attempts to generate stable METTL16-null TNBC cell lines were unsuccessful due to profound loss of cell viability. These findings indicate that METTL16 is essential for maintaining TNBC cell survival and is required for fundamental cellular processes in these cells. Together, these results demonstrate that METTL16 not only promotes resistance to taxanes but is also indispensable for TNBC cell survival. In contrast, knockout of METTL16 did not markedly impair the survival of non-cancerous human mammary epithelial (hTERT-HME1) cells (Fig. 8 F,G). Although CRISPR-Cas9–mediated depletion of METTL16 resulted in a modest (∼30%) reduction in HME cell survival compared with control cells (Fig. 8 H), this partial effect is consistent with a fundamental role for METTL16 in maintaining normal cellular homeostasis (Doxtader et al. 2018). Importantly, the degree of dependency on METTL16 was substantially greater in TNBC cells, highlighting a cancer-selective reliance on METTL16 for survival.

**Fig 8.**
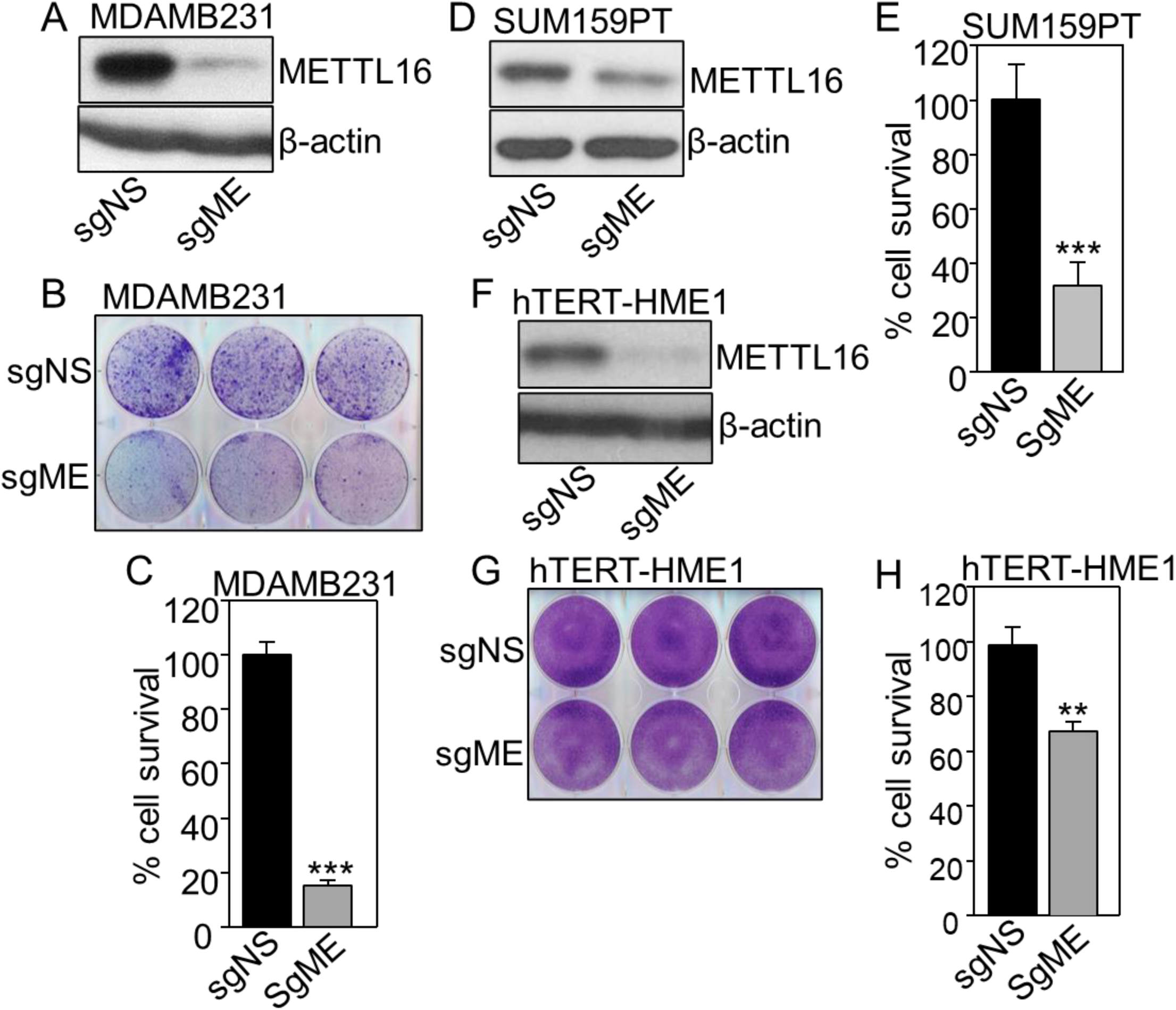
METTL16 is essential for the survival of paclitaxel-sensitive TNBC cells but not a non-tumorigenic mammary epithelial cell line. **A.** Western analysis of METTL16 in MDAMB231 cells transduced with a CRISPR-Cas9 lentivirus expressing METTL16-targeting sgRNA, compared to non-targeting control. β-actin was used as a loading control. sgNS: non-targeting control and sgME: METTL16-targeting sgRNA. **B.** Images from colony formation assays. MDAMB231 cells were seeded after down-regulation of METTL16 and cultured for 11 days. Colonies were fixed and stained with crystal violet. **C.** Cell survival assays were performed with MDAMB231 cells expressing sgNS or sgME. Cells were plated in 96-well plates at a density of 4,000 cells per well, and after 13 days, cell viability was assessed using the Alamar Blue reagent. **D.** Western analysis of METTL16 in SUM159PT cells transduced with CRISPR-Cas9 lentivirus expressing METTL16-targeting sgRNA compared to non-targeting control. β-actin was used as a loading control. **E.** Cell survival assays were performed in SUM159PT cells expressing sgNS or sgME. Cells were plated in 96-well plates at a density of 500 cells per well, and after 7 days, cell viability was assessed using the Alamar Blue reagent. **F.** Western analysis of METTL16 in HME cells transduced with a CRISPR-Cas9 lentivirus expressing METTL16-targeting sgRNA compared to non-targeting control. β-actin was used as a loading control. **G.** Images from colony formation assays. HME cells were seeded after down-regulation of METTL16 and cultured for 5 days. Colonies were fixed and stained with crystal violet. **H.** Cell survival assays were performed in HME cells expressing sgNS or sgME. Cells were plated in 48-well plates at a density of 10,000 cells per well, and after 13 days cell viability was assessed using 1 M NaOH. Results are represented by means ± SD. Data were analyzed using Student’s t-test. **, P< 0.01; ***, P< 0.001.

## Discussion

In this study, we identified METTL16 as a central epi-transcriptomic regulator that links m^6^A-dependent translational control to taxane resistance and survival in TNBC. Using complementary genetic, biochemical, translational, and *in vivo* approaches, we demonstrate that METTL16 promotes resistance by methylating ABCB1 mRNA, thereby enhancing its translation and increasing drug efflux. While ABCB1 overexpression has long been recognized as a hallmark of multidrug resistance, the upstream epi-transcriptomic mechanisms governing its dysregulation are not well defined in TNBC. Our findings identify METTL16 as a previously unrecognized translational regulator of ABCB1, expanding the emerging concept that m^6^A-dependent RNA modification contributes to chemotherapy resistance.

Mechanistically, our data identify ABCB1 as a functionally critical METTL16 target. We show that METTL16 binds ABCB1 mRNA and promotes m^6^A deposition across multiple regions of the transcript. Although the magnitude of ABCB1 enrichment in RNA-IP was lower than that observed for MAT2A, its marked enrichment above IgG control supports ABCB1 as a noncanonical METTL16-associated transcript with functional relevance to taxane resistance. Importantly, this regulation occurs without a corresponding increase in ABCB1 mRNA abundance, indicating that METTL16 controls ABCB1 expression primarily at the level of translation rather than transcription or mRNA stability. Consistent with this model, polysome profiling revealed that METTL16 enhances ribosome loading onto ABCB1 mRNA, shifting transcripts from monosomal to polysomal fractions and increasing translational efficiency (Ambar-Martinez and Young-Baird 2025). m^6^A deposition can enhance mRNA stability or translation through recruitment of m^6^A reader proteins (Jiang et al. 2021), thereby increasing steady-state ABCB1 expression and promoting drug efflux capacity. Future studies will determine which m^6^A reader protein/s mediate METTL16-dependent translational enhancement of ABCB1 and define the precise molecular mechanisms linking m^6^A deposition to increased drug efflux capacity. This translational enhancement was dependent on METTL16 catalytic activity, as expression of a catalytically inactive mutant reduced ABCB1 polysome association.

The requirement for METTL16 catalytic activity is particularly significant. While METTL16 has been reported to possess non-catalytic functions, including RNA binding and structural roles (Su et al. 2022), our results demonstrate that m^6^A deposition is essential for its resistance function. Catalytic inactivation of METTL16 abolished its ability to confer taxane resistance and partially suppressed resistance in PacR cells, highlighting m^6^A methylation as the key functional output driving this phenotype. Notably, although METTL3-METTL14 has been widely implicated in cancer-associated m^6^A regulation (Yuan et al. 2025), our findings identify METTL16 as a distinct and context-specific regulator of translational control in TNBC. Unlike METTL3, which broadly influences transcript stability and translation across the transcriptome, METTL16 appears to regulate a more selective subset of RNAs, suggesting that therapeutic targeting of METTL16 may achieve greater specificity with reduced global disruption of RNA metabolism. These findings underscore the importance of the enzymatic activity of METTL16 as a therapeutic target.

Our Flutax accumulation studies indicate that METTL16 regulates taxane resistance by controlling intracellular drug levels. By increasing ABCB1-mediated efflux, METTL16 lowers paclitaxel concentrations inside the cell, thereby limiting effective mitotic arrest and activation of apoptosis (Khing et al. 2021). Calcein-AM efflux assays further support this mechanism by confirming active transporter function in resistant cells. When METTL16 is depleted, intracellular paclitaxel retention increases, restoring the ability of the drug to trigger apoptotic cell death. These findings suggest that METTL16 promotes resistance, at least in part, by regulating ABCB1 translation and intracellular drug accumulation.

Our clinical correlation analyses further support the relevance of the METTL16-ABCB1 axis in human disease. The positive association between METTL16 and ABCB1 expression in TNBC patient cohorts, particularly within the mesenchymal-like immune-altered subtype, suggests that this pathway may be especially important in aggressive, therapy-refractory tumors (Jézéquel et al. 2024). Notably, the TNBC models used in our mechanistic studies belong to this mesenchymal/stem-like category (Huang et al. 2024), reinforcing the translational relevance of our findings. From a therapeutic perspective, our work highlights METTL16 as an attractive upstream target for overcoming drug resistance. Direct inhibition of ABCB1 has proven challenging due to toxicity, redundancy among ABC transporters, and compensatory resistance mechanisms (Tamaki et al. 2011). By contrast, targeting METTL16 disrupts ABCB1 expression indirectly while simultaneously impairing TNBC cell survival. Our Vivo-Morpholino studies demonstrate that pharmacologic inhibition of METTL16 translation significantly reduces viability of PacR TNBC cells and suppresses tumor growth *in vivo*, supporting the feasibility of this strategy and its potential therapeutic selectivity. These findings expand the functional landscape of m^6^A RNA methyltransferases in cancer and highlight METTL16 as a promising therapeutic target for overcoming taxane resistance in aggressive breast cancers.

A key insight from our work is that METTL16 functions as both a resistance driver and a survival dependency in TNBC cells. Genetic down-regulation of METTL16 resulted in profound loss of viability across multiple TNBC models, whereas non-cancerous mammary epithelial cells were only modestly affected. This differential dependency suggests that TNBC cells rely on METTL16-regulated RNA processing pathways to maintain cellular homeostasis under stress conditions such as chemotherapy exposure. The dual role of METTL16 as both a resistance mediator and a survival dependency creates a potential therapeutic window in which targeting METTL16 simultaneously impairs drug efflux and compromises tumor cell viability, thereby limiting the capacity of resistant cells to adapt.

Several questions remain for future investigation. While ABCB1 represents a major downstream effector of METTL16-mediated resistance, it is likely not the only relevant target. METTL16 has been reported to regulate a limited but functionally important subset of RNAs (Brown et al. 2016; Mermoud 2022) and investigating the full METTL16-dependent translational landscape in TNBC will be important. In addition, the mechanisms driving METTL16 upregulation in paclitaxel-resistant cells remain to be elucidated.

## Authors’ Contributions

Conception and design: S.D

Development of methodology: E.G.H.B, J.A.C, D.J.L, A.A, S.D.

Acquisition of data: E.G.H.B, J.A.C, D.J.L, A.A, A.B, S.D.

Analysis and interpretation of data: E.G.H.B, J.A.C, D.J.L, A.A.K, S.D.

Writing, review, and/or revision of the manuscript: E.G.H.B, J.A.C, H.G, A.A.K, G.R.S, S.D.

Administrative, technical, or material support: E.G.H.B, Y.P.

Study supervision: G.R.S, S.D.

## Acknowledgments

This research was supported by a Catalyst Pilot grant from Cleveland Clinic (to SD), PPG, NCI (P01CA272161; to GRS), and in part by NIGMS (R01GM128981; to AAK). We are extremely thankful to Dr. Ruth Keri, Cleveland Clinic for providing paclitaxel resistant cell lines. We would like to thank Drs. Rui Su and Jianjun Chen (Beckman Research Institute of City of Hope, Monrovia, CA) for the METTL16 sgRNA constructs. We thank the Cleveland Clinic Flow Cytometry Core for excellent technical support.

## References

Abd El-Aziz YS, Spillane AJ, Jansson PJ, et al. 2021. Role of ABCB1 in mediating chemoresistance of triple-negative breast cancers. Biosci Rep. 41:BSR20204092.

Aartsma-Rus A, Krieg AM. 2017. FDA approves eteplirsen for Duchenne muscular dystrophy. Nucleic Acid Ther. 27:1.

Abu Samaan TM, Samec M, Liskova A, et al. 2019. Paclitaxel mechanistic and clinical effects in breast cancer. Biomolecules. 9:789.

Alalawy AI. 2024. Key genes and molecular mechanisms related to paclitaxel resistance. Cancer Cell Int. 24:244.

Ambar-Martinez R, Young-Baird SK. 2025. Polysome profiling as a tool to study translation. Mol Biol Cell. 36:mr2.

Bergonzini C, Gregori A, Hagens TMS, et al. 2024. ABCB1 overexpression via locus amplification drives paclitaxel resistance. J Exp Clin Cancer Res. 43:4.

Brown JA, Kinzig CG, DeGregorio SJ, et al. 2016. Methyltransferase-like protein 16 binds the 3′-terminal triple helix of MALAT1 long noncoding RNA. Proc Natl Acad Sci USA. 113:14013.

Cerneckis J, Ming GL, Song H, et al. 2024. The rise of epitranscriptomics: recent developments and future directions. Trends Pharmacol Sci. 45:24–38.

Chen Z, Liu Y, Lyu M, et al. 2025. Classifications of triple-negative breast cancer: insights and current therapeutic approaches. Cell Biosci. 15:13.

De S, Cipriano R, Jackson MW, et al. 2009. Overexpression of kinesins mediates docetaxel resistance in breast cancer cells. Cancer Res. 69:8035–8042.

De S, Dermawan JK, Stark GR. 2014. EGF receptor uses SOS1 to drive constitutive activation of NFκB in cancer cells. Proc Natl Acad Sci USA. 111:11721–11726.

De S, Lindner DJ, Coleman CJ, et al. 2018. The FACT inhibitor CBL0137 synergizes with cisplatin in small-cell lung cancer by increasing NOTCH1 expression and targeting tumor-initiating cells. Cancer Res. 78:2396–2406.

Doxtader KA, Wang P, Scarborough AM, et al. 2018. Structural basis for regulation of METTL16, an S-adenosylmethionine homeostasis factor. Mol Cell. 71:1001–1011.

Ferguson DP, Schmitt EE, Lightfoot JT. 2013. Vivo-morpholinos induced transient knockdown of physical activity related proteins. PLoS One. 8:e61472.

Flaherty JN, Sivasudhan E, Tegowski M, et al. 2025. Catalytic efficiency of METTL16 regulates cellular SAM homeostasis. Cell Rep. 44:115966.

Frayde-Gómez H, Chimal-Vega B, Pulido-Capiz A, et al. 2026. The translation factor eIF4E mediates doxorubicin resistance. Sci Rep. 16:1805.

Gruber AR, Lorenz R, Bernhart SH, et al. 2008. The Vienna RNA Websuite. Nucleic Acids Res. 36:W70–W74.

Győrffy B. 2021. Survival analysis across the transcriptome identifies biomarkers in breast cancer. Comput Struct Biotechnol J. 19:4101–4109.

Huang P, Zhang X, Prabhu JS, et al. 2024. Therapeutic vulnerabilities in triple negative breast cancer. Biomed Pharmacother. 174:116584.

Jézéquel P, Lasla H, Gouraud W, et al. 2024. Mesenchymal-like immune-altered TNBC subtype. Breast Cancer. 31:825–840.

Jiang X, Liu B, Nie Z, et al. 2021. Role of m6A modification in biological functions and diseases. Signal Transduct Target Ther. 6:74.

Khing TM, Choi WS, Kim DM, et al. 2021. The effect of paclitaxel on apoptosis, autophagy and mitotic catastrophe in AGS cells. Sci Rep. 11:23490.

Liu N, Zhou KI, Parisien M, et al. 2017. N6-methyladenosine alters RNA structure to regulate binding of a low-complexity protein. Nucleic Acids Res. 45:6051–6063.

Lu T, Jackson MW, Singhi AD, et al. 2009. Validation-based insertional mutagenesis identifies FBXL11 as a regulator of NFκB. Proc Natl Acad Sci USA. 106:16339–16344.

Maloney SM, Hoover CA, Morejon-Lasso LV, et al. 2020. Mechanisms of taxane resistance. Cancers (Basel). 12:3323.

Matsui A, Ihara T, Suda H, et al. 2013. Gene amplification: mechanisms and involvement in cancer. Biomol Concepts. 4:567–582.

Mermoud JE. 2022. The role of the m6A RNA methyltransferase METTL16 in gene expression and SAM homeostasis. Genes (Basel). 13:2312.

Moreno-Aspitia A, Perez EA. 2009. Treatment options for breast cancer resistant to anthracycline and taxane. Mayo Clin Proc. 84:533–545.

Moulton JD, Jiang S. 2009. Gene knockdowns in adult animals: PPMOs and Vivo-Morpholinos. Molecules. 14:1304–1323.

Obidiro O, Battogtokh G, Akala EO. 2023. Triple negative breast cancer treatment options and limitations: future outlook. Pharmaceutics. 15:1796.

Parker BJ, Moltke I, Roth A, et al. 2011. New families of human regulatory RNA structures. Genome Res. 21:1929–1943.

Pendleton KE, Chen B, Liu K, et al. 2017. The U6 snRNA m6A methyltransferase METTL16 regulates SAM synthetase intron retention. Cell. 169:824–835.

Piska K, Koczurkiewicz-Adamczyk P, Jamrozik M, et al. 2023. ABCB1-dependent efflux of anthracyclines and metabolites. Xenobiotica. 53:507–514.

Pucci G, Minafra L, Bravatà V, et al. 2024. Glut-3 gene knockdown as a strategy to overcome glioblastoma radioresistance. Int J Mol Sci. 25:2079.

Roberts MS, Sahni JM, Schrock MS, et al. 2020. LIN9 and NEK2 are core regulators of mitotic fidelity that can be therapeutically targeted to overcome taxane resistance. Cancer Res. 80:1693–1706.

Roshmi RR, Yokota T. 2023. Viltolarsen: from preclinical studies to FDA approval. Methods Mol Biol. 2587:31–41.

Ruszkowska A. 2021. METTL16: current insights into structure and function. Int J Mol Sci. 22:2176.

Ruszkowska A, Ruszkowski M, Dauter Z, et al. 2018. Structural insights into the RNA methyltransferase domain of METTL16. Sci Rep. 8:5311.

Satterwhite ER, Mansfield KD. 2022. RNA methyltransferase METTL16: targets and function. WIREs RNA. 13:e1681.

Su R, Dong L, Li Y, et al. 2022. METTL16 exerts an m6A-independent function to facilitate translation and tumorigenesis. Nat Cell Biol. 24:205–216.

Tamaki A, Ierano C, Szakacs G, et al. 2011. The controversial role of ABC transporters in clinical oncology. Essays Biochem. 50:209.

Tan MH, De S, Bebek G, et al. 2012. Specific kinesin expression profiles associated with taxane resistance. Breast Cancer Res Treat. 131:849–858.

Tian Y, Lei Y, Wang Y, et al. 2023. Mechanism of multidrug resistance mediated by P-glycoprotein. Int J Oncol. 63:119.

Wang H, Xu B, Shi J. 2020. METTL3 promotes breast cancer progression via Bcl-2. Gene. 722:144076.

Wang Q, Jiang X, Yuan Y, et al. 2025. METTL16 emerges as a pivotal epitranscriptomic regulator linking RNA modification, tumor progression, and immune modulation. Front Immunol. 16:1706971.

Xu J, Gan C, Yu S, et al. 2024. Analysis of immune resistance mechanisms in TNBC: dual effects inside and outside the tumor. Clin Breast Cancer. 24:e91–e102.

Xuan JJ, Sun WJ, Lin PH, et al. 2018. RMBase v2.0: RNA modification database. Nucleic Acids Res. 46:D327–D334.

Ye F, Wu J, Zhang F. 2023. METTL16 epigenetically enhances GPX4 expression via m6A modification to promote breast cancer progression. Biochem Biophys Res Commun. 638:1–6.

Yi D, Wang R, Shi X, et al. 2020. METTL14 promotes migration and invasion of breast cancer cells. Oncol Rep. 43:1375–1386.

Yoshinaga M, Han K, Morgens DW, et al. 2022. METTL16 enables erythropoiesis through genome integrity. Nat Commun. 13:6435.

Yuan F, Zhang W, Xia Y, et al. 2025. Role of METTL3 in cancer and therapeutic potential. Eur J Med Res. 30:1017.

Zhang R, Zhang Y, Guo F, et al. 2022. Knockdown of METTL16 disrupts learning and memory by reducing MAT2A mRNA stability. Cell Death Discov. 8:432.

Zhou Y, Zeng P, Li YH, et al. 2016. SRAMP: prediction of mammalian m6A sites. Nucleic Acids Res. 44:e91.

