## Supplementary Figures for "METTL16 promotes taxane resistance in Triple-Negative Breast Cancer through m^6^A-dependent translational upregulation of ABCB1"

Suppl. Fig. 1

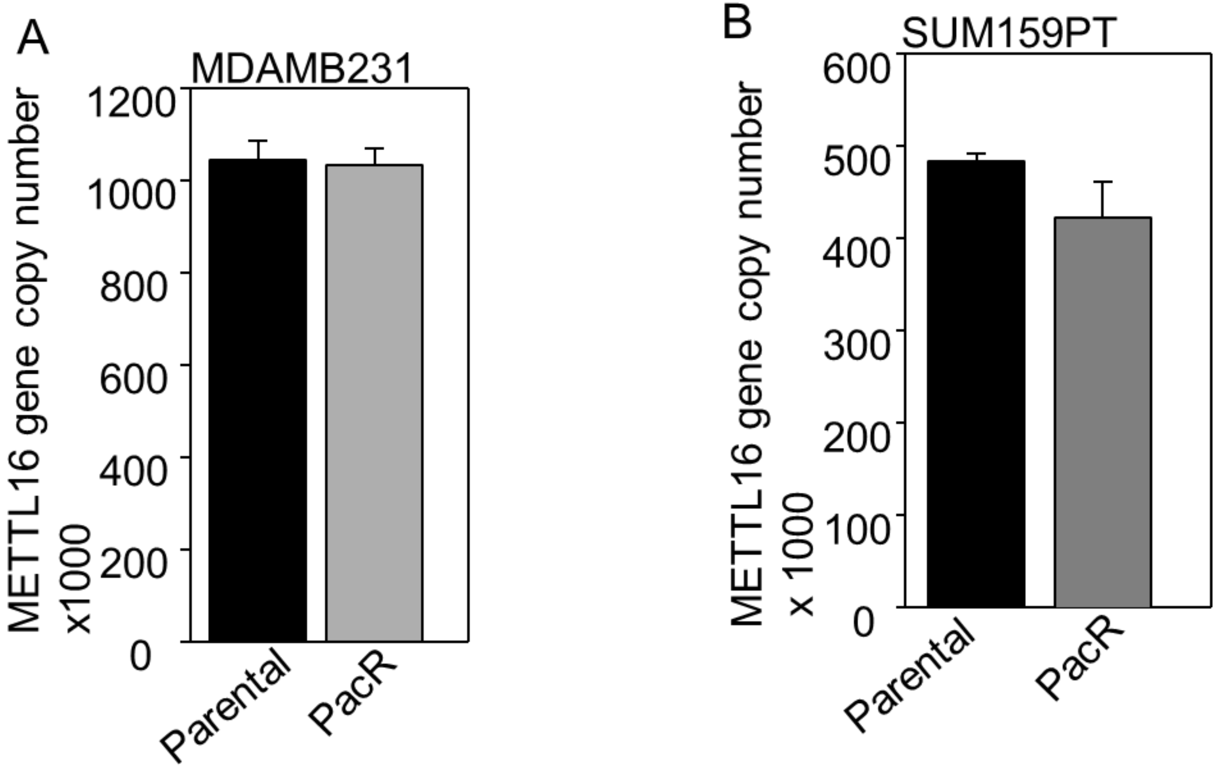

A
MDAMB231
METTL16 gene copy number x1000
1200
1000
800
600
400
200
0
Parental
PacR
B
SUM159PT
METTL16 gene copy number x 1000
600
500
400
300
200
100
0
Parental
PacR

Suppl. Fig. 2

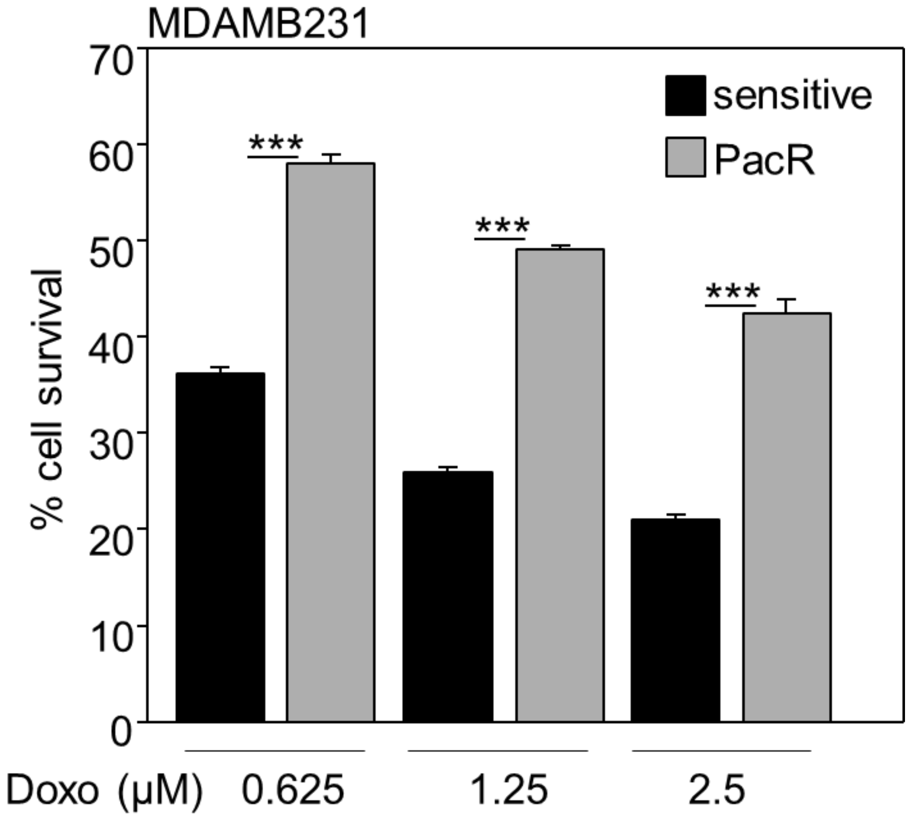
MDAMB231
sensitive
PacR
***
***
***
% cell survival
0
10
20
30
40
50
60
70
Doxo (μM)
0.625
1.25
2.5

Suppl. Fig. 3

A

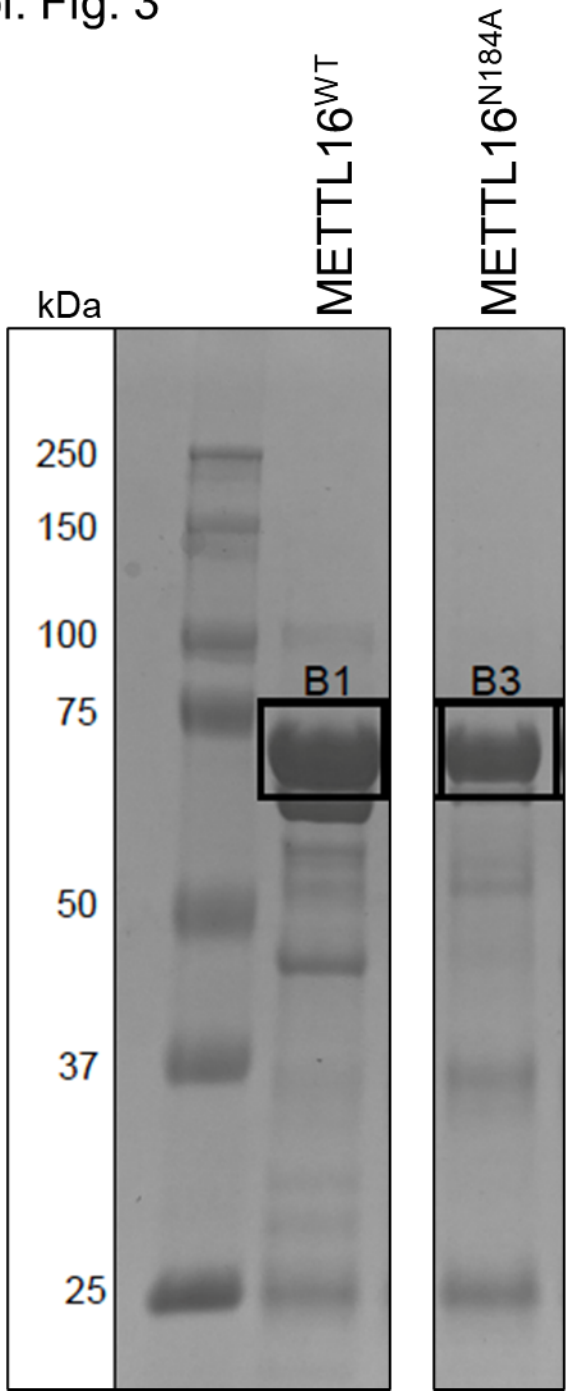


B

METTL16$^{WT}$ Band 1 Identified Tryptic Peptides (87% coverage)

```
1   MGHHHHHHSS GLEVLFQGPM ALSKSMHARN RYKDKPPDFA YLASKYPDFK QHVQINLNGR VSLNFKDPEA VRALTCTLLR
81  EDFGLSIDIP LERLIPTVPL RLNYIHWVED LIGHQDSDKS TLRRGIDIGT GASCIYPLLG ATLNGWYFLA TEVDDMCFNY
161 AKKNVEQNNL SDLIKVVKVP QKTLLMDALK EESEIIYDFC MCNPPFFANQ LEAKGVNSRN PRRPPPSSVN TGGITEIMAE
241 GGELEFVKRI IHDSLQLKKR LRWYSCMLGK KCSLAPLKEE LRIQGVPKVT YTEFCQGRTM RWALAWSFYD DVTVPSPPSK
321 RRKLEKPRKP ITFVVLASVM KELSLKASPL RSETAEGIVV VTTWIEKILT DLKVQHKRVP CGKEEVSLFL TAIENSWIHL
401 RRKKRERVRQ LREVPRAPED VIQALEEKKP TPKESGNSQE LARGPQERTP CGPALREGEA AAVEGPCPSQ ESLSQEENPE
481 PTEDERSEEK GGVEVLESCQ GSSNGAQDQE ASEQFGSPVA ERGKRLPGVA GQYLFKCLIN VKKEVDDALV EMHWVEGQNR
561 DLMNQLCTYI RNQIFRLVAV N
```

METTL16$^{N184}$ Band 3 Identified Tryptic Peptides (77% coverage)

```
1   MGHHHHHHSS GLEVLFQGPM ALSKSMHARN RYKDKPPDFA YLASKYPDFK QHVQINLNGR VSLNFKDPEA VRALTCTLLR
81  EDFGLSIDIP LERLIPTVPL RLNYIHWVED LIGHQDSDKS TLRRGIDIGT GASCIYPLLG ATLNGWYFLA TEVDDMCFNY
161 AKKNVEQNNL SDLIKVVKVP QKTLLMDALK EESEIIYDFC MCNPPFFANQ LEAKGVNSRN PRRPPPSSVN TGGITEIMAE
241 GGELEFVKRI IHDSLQLKKR LRWYSCMLGK KCSLAPLKEE LRIQGVPKVT YTEFCQGRTM RWALAWSFYD DVTVPSPPSK
321 RRKLEKPRKP ITFVVLASVM KELSLKASPL RSETAEGIVV VTTWIEKILT DLKVQHKRVP CGKEEVSLFL TAIENSWIHL
401 RRKKRERVRQ LREVPRAPED VIQALEEKKP TPKESGNSQE LARGPQERTP CGPALREGEA AAVEGPCPSQ ESLSQEENPE
481 PTEDERSEEK GGVEVLESCQ GSSNGAQDQE ASEQFGSPVA ERGKRLPGVA GQYLFKCLIN VKKEVDDALV EMHWVEGQNR
561 DLMNQLCTYI RNQIFRLVAV N
```

Suppl. Fig. 4

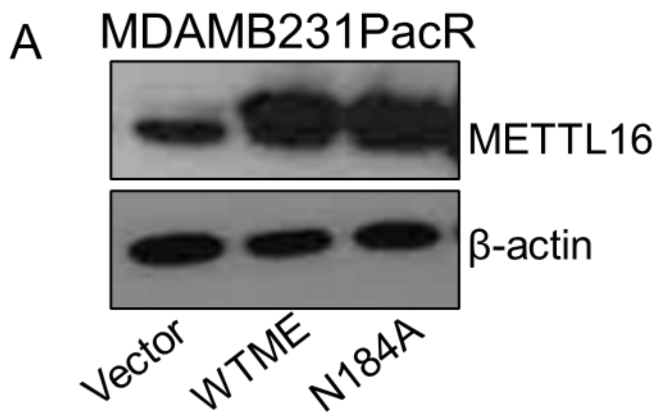
A
MDAMB231PacR
METTL16
β-actin
Vector
WTME
N184A

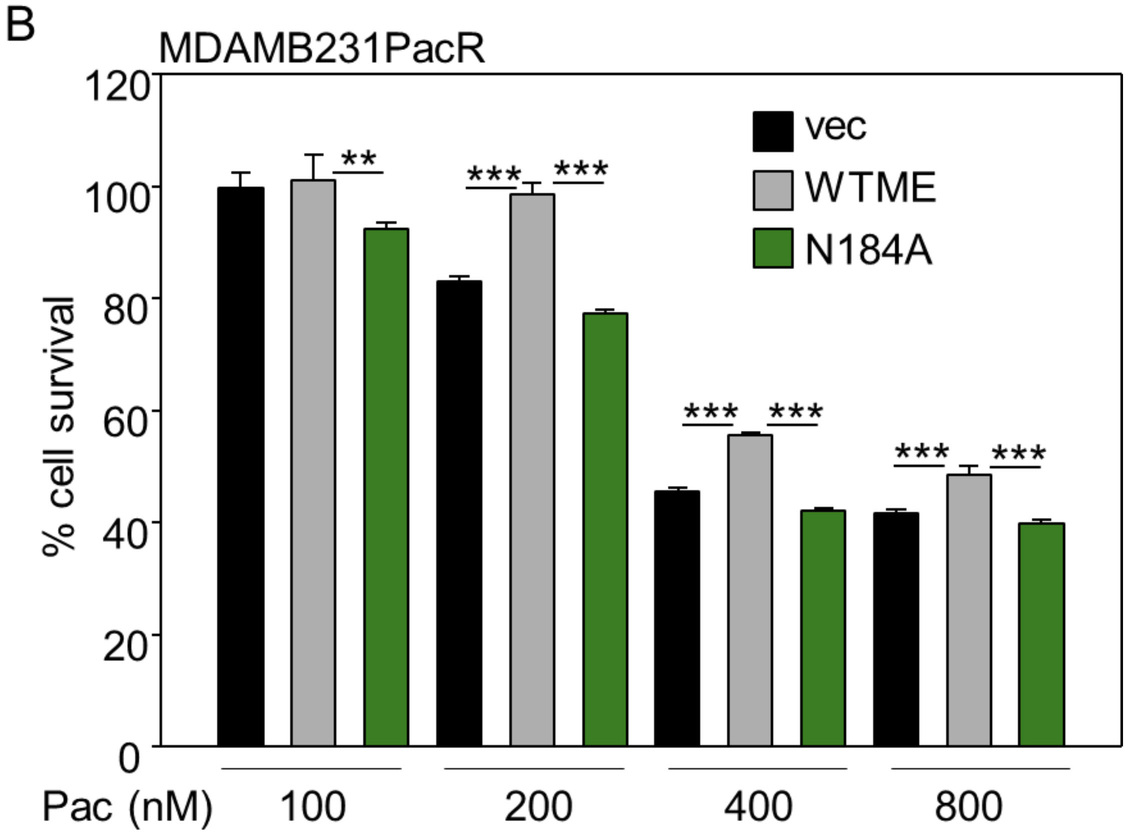
B
MDAMB231PacR
% cell survival
120
100
80
60
40
20
0
vec
WTME
N184A
**
***
Pac (nM)
100
200
400
800

Suppl. Fig. 5

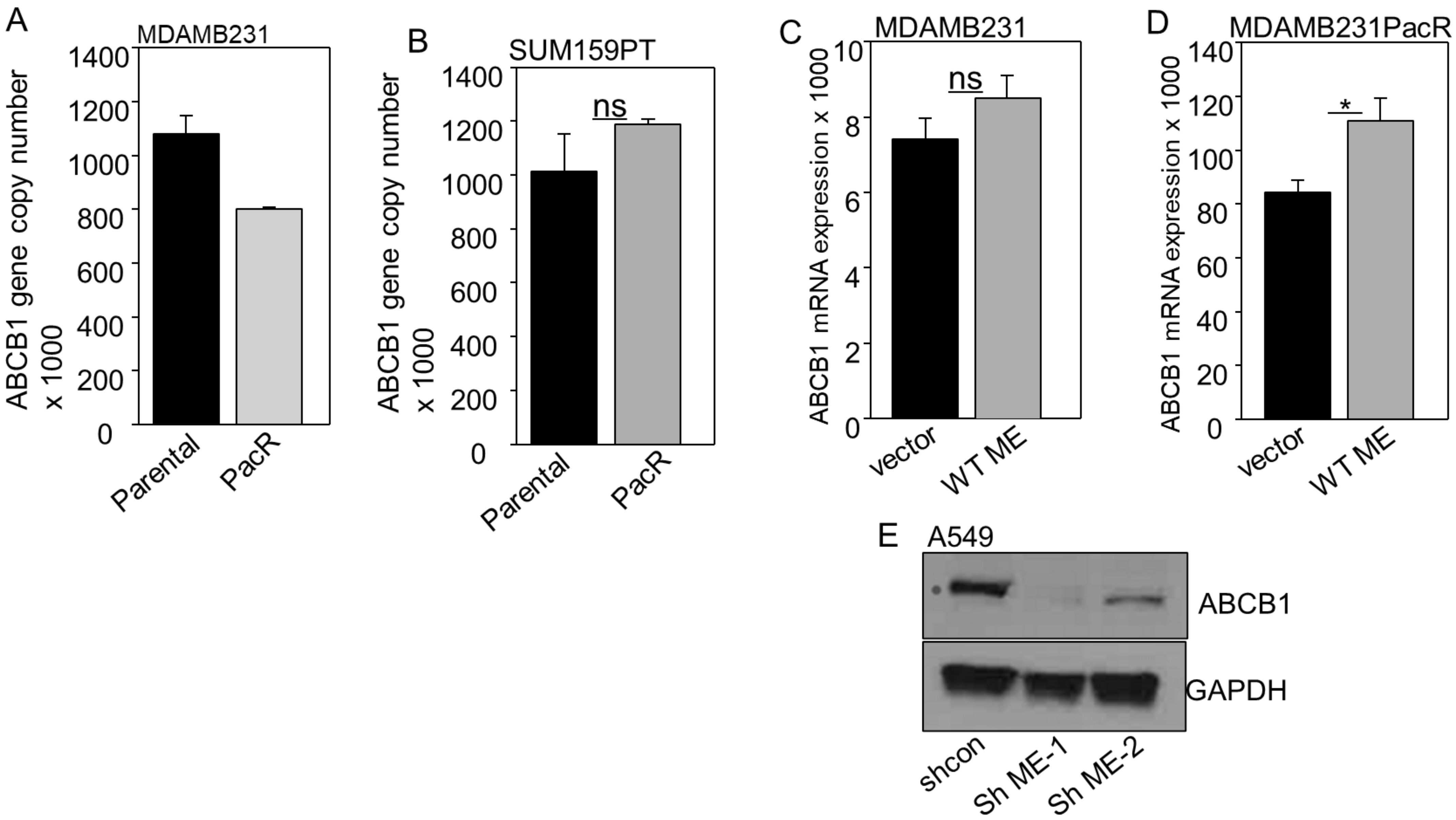
A
MDAMB231
ABCB1 gene copy number x 1000
1400
1200
1000
800
600
400
200
0
Parental
PacR
B
SUM159PT
ns
ABCB1 gene copy number x 1000
1400
1200
1000
800
600
400
200
0
Parental
PacR
C
MDAMB231
ns
ABCB1 mRNA expression x 1000
10
8
6
4
2
0
vector
WT ME
D
MDAMB231PacR
*
ABCB1 mRNA expression x 1000
140
120
100
80
60
40
20
0
vector
WT ME
E
A549
ABCB1
GAPDH
shcon
Sh ME-1
Sh ME-2

Suppl. Fig. 6

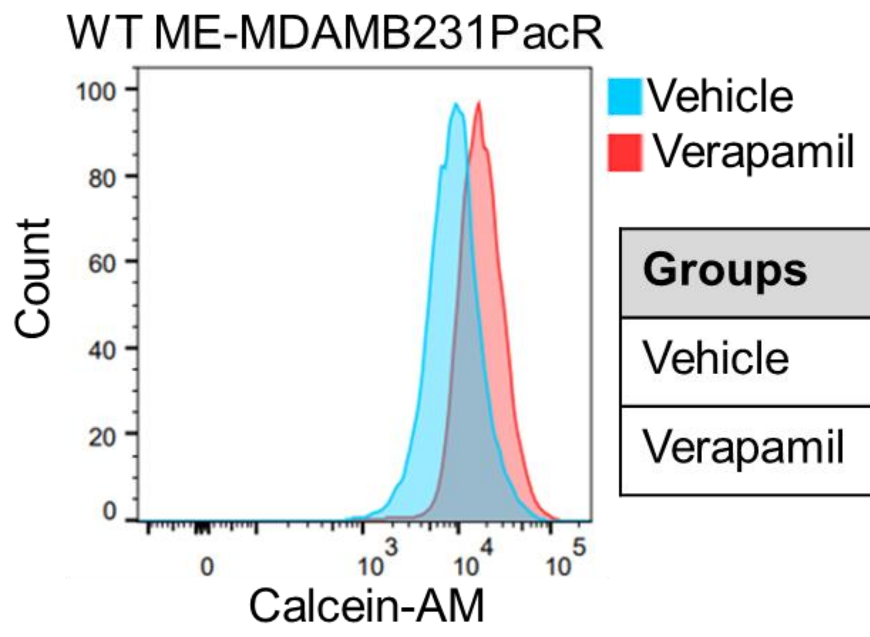


| Groups | Median |
|---|---|
| Vehicle | 9086 |
| Verapamil | 17403 |

Suppl. Fig. 7

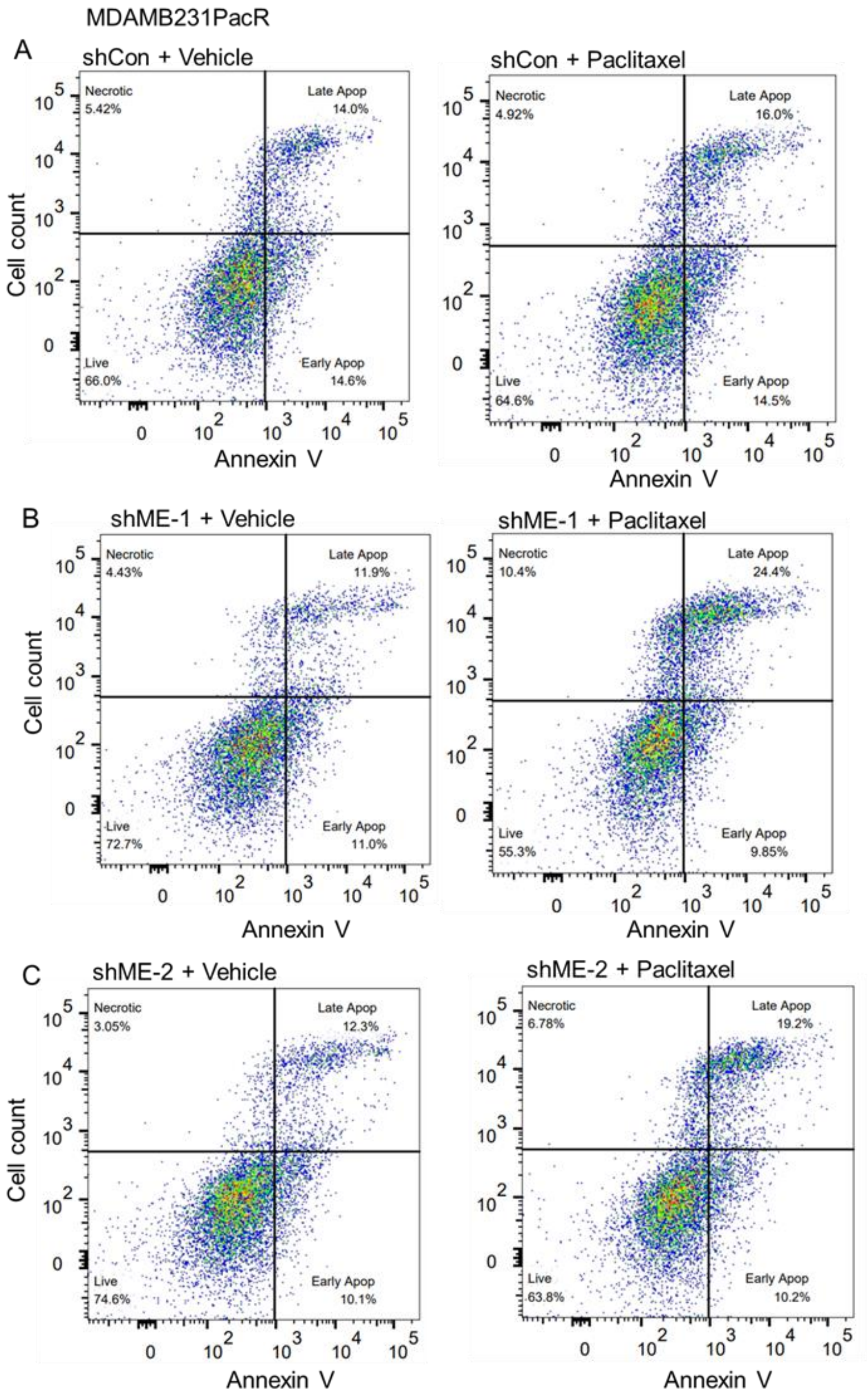

MDAMB231PacR
A
shCon + Vehicle
Necrotic 5.42%
Late Apop 14.0%
Live 66.0%
Early Apop 14.6%
shCon + Paclitaxel
Necrotic 4.92%
Late Apop 16.0%
Live 64.6%
Early Apop 14.5%
B
shME-1 + Vehicle
Necrotic 4.43%
Late Apop 11.9%
Live 72.7%
Early Apop 11.0%
shME-1 + Paclitaxel
Necrotic 10.4%
Late Apop 24.4%
Live 55.3%
Early Apop 9.85%
C
shME-2 + Vehicle
Necrotic 3.05%
Late Apop 12.3%
Live 74.6%
Early Apop 10.1%
shME-2 + Paclitaxel
Necrotic 6.78%
Late Apop 19.2%
Live 63.8%
Early Apop 10.2%
Cell count
Annexin V

Suppl. Fig. 8

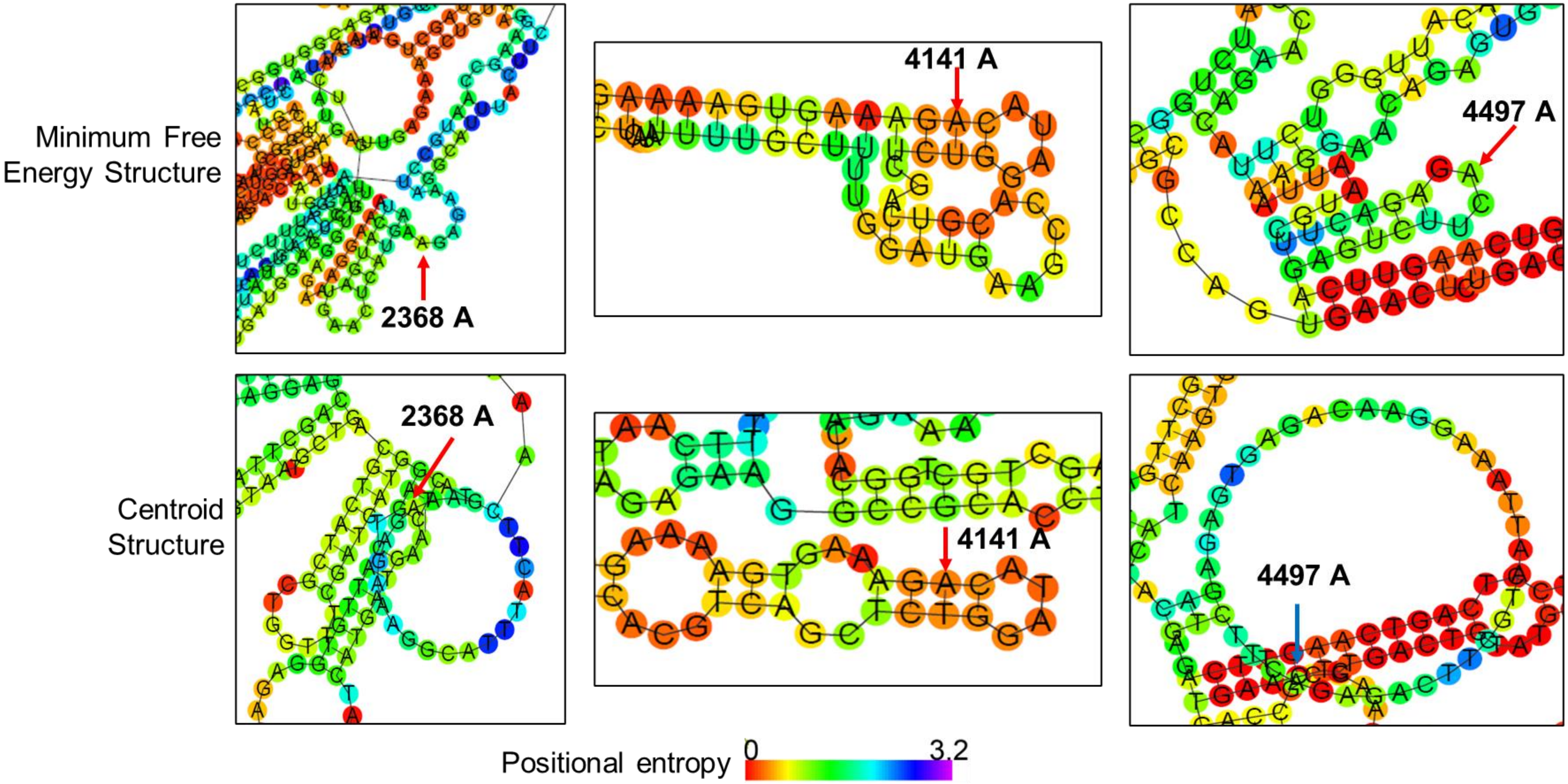
Minimum Free Energy Structure
Centroid Structure
2368 A
4141 A
4497 A
2368 A
4141 A
4497 A
Positional entropy
0
3.2

Suppl. Fig. 9

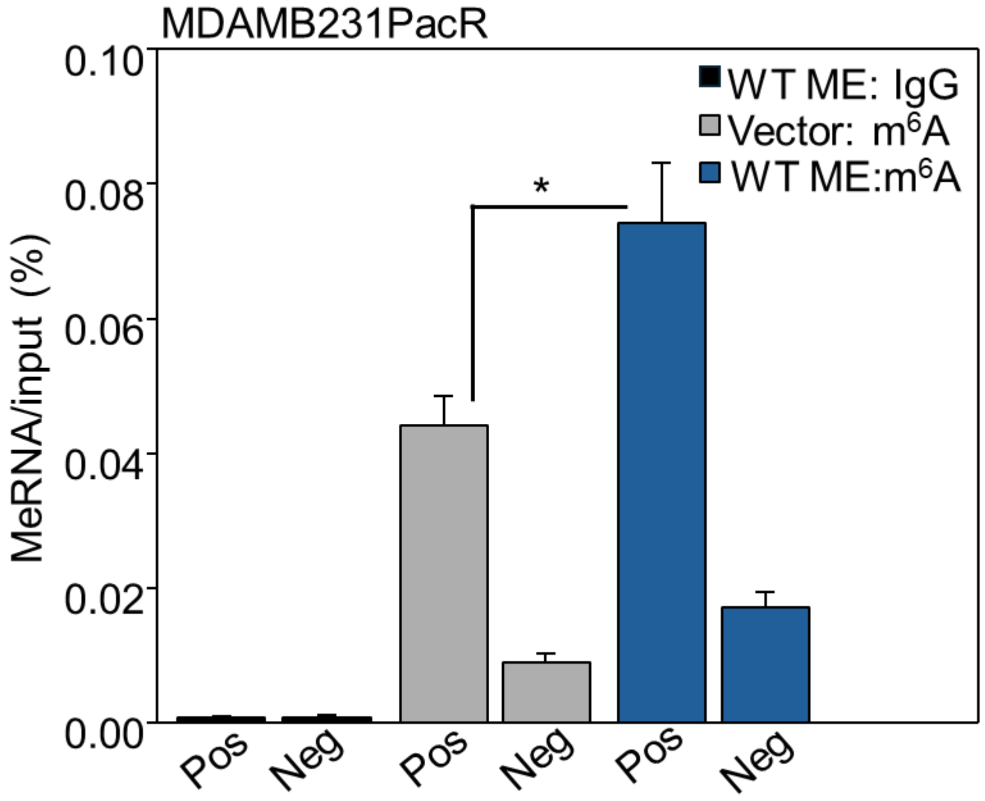

MDAMB231PacR
MeRNA/input (%)
0.10
0.08
0.06
0.04
0.02
0.00
WT ME: IgG
Vector: m6A
WT ME:m6A
*
Pos
Neg
Pos
Neg
Pos
Neg

Suppl. Fig. 10

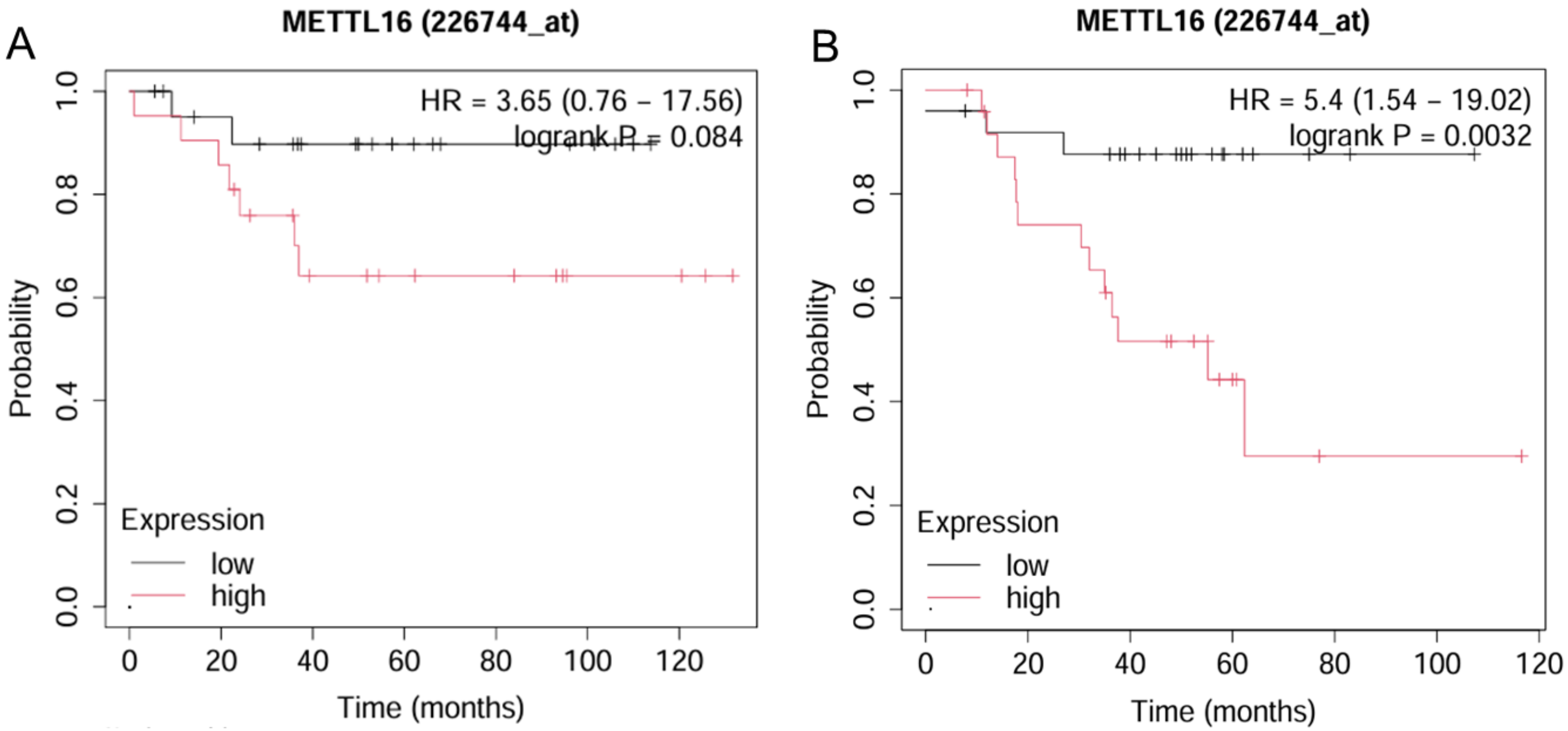
A
METTL16 (226744_at)
HR = 3.65 (0.76 – 17.56)
logrank P = 0.084
Probability
Time (months)
Expression
low
high
B
METTL16 (226744_at)
HR = 5.4 (1.54 – 19.02)
logrank P = 0.0032
Probability
Time (months)
Expression
low
high

Suppl. Fig. 11

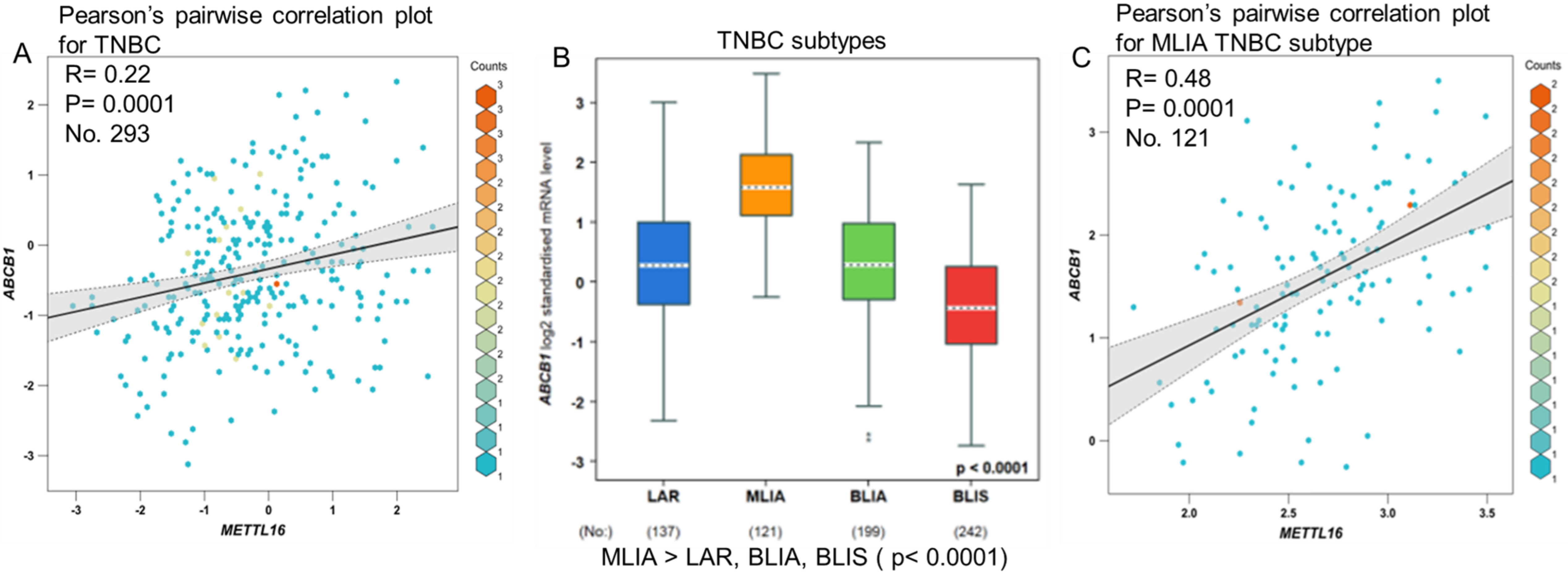
A
Pearson's pairwise correlation plot for TNBC
R= 0.22
P= 0.0001
No. 293
Counts
ABCB1
METTL16
B
TNBC subtypes
ABCB1 log2 standardised mRNA level
p < 0.0001
LAR
MLIA
BLIA
BLIS
(No:)
(137)
(121)
(199)
(242)
MLIA > LAR, BLIA, BLIS ( p< 0.0001)
C
Pearson's pairwise correlation plot for MLIA TNBC subtype
R= 0.48
P= 0.0001
No. 121
Counts
ABCB1
METTL16
