## Supplemental Methods for "METTL16 promotes taxane resistance in Triple-Negative Breast Cancer through m^6^A-dependent translational upregulation of ABCB1"

**Recombinant METTL16 Purification**

Full-length human METTL16 (UniProt: Q86W5-1; Met1—Asn562) with an N-terminal, HRV3C-cleavable His6 tag was ordered as an *E. coli* codon optimized construct in pET28A+ (Twist Biosciences). The N184A mutation, located within the conserved catalytic motif of the methyltransferase domain, was generated by site-directed mutagenesis (Agilent QuikChange) and confirmed by Sanger sequencing. Both $METTL16^{WT}$ and $METTL16^{N184A}$ were purified essentially as described by Ruszkowska *et al.* (2018). Constructs were transformed in BL21 Gold cells (Agilent), grown to OD = 0.8 in LB, dropped to 18 °C, induced with 0.5 mM IPTG, and expressed overnight. Cell pellets were broken by sonication in Base Buffer (50 mM HEPES pH = 7.5, 500 mM NaCl, 5% glycerol, 0.5 mM TCEP) supplemented with cOmplete™ protease inhibitors (Roche) and benzonase (in-house purified), clarified by centrifugation, purified by IMAC over TALON® resin (Takara), and polished on a Superdex 200 16/600 column (GE) into Gel Filtration Buffer (25 mM HEPES pH = 7.5, 100 mM KCl, 50 mM NaCl, 1 mM TCEP). The monodisperse, monomeric peak was concentrated to 1-2 mg/mL (Amicon centrifugal concentrator) and snap-frozen prior to long-term storage at -80 °C. Protein identity was validated by in-gel tryptic digest MSMS and purity was assessed by SDS-PAGE.

**Calcein-AM Efflux Assay**

ABCB1 transport activity was assessed using a Calcein-AM efflux assay. WT METTL16 overexpressing MDA-MB-231-PacR cells were grown to 70-80% confluency and pre-treated with verapamil (5 µM; Sigma Aldrich, cat. # 676777) or vehicle control for 30 minutes at 37°C. Calcein acetoxymethyl ester (Calcein-AM; 0.25 µM; Invitrogen, cat. # C1430) was then added directly to the medium, and cells were incubated for an additional 30 minutes at 37°C in the continued presence or absence of verapamil. Following incubation, cells were washed twice with PBS, detached by brief trypsinization, and resuspended in PBS for flow cytometric analysis. Intracellular fluorescence was measured using a BD FACSymphony A5 SE (5-laser configuration; BD Biosciences, San Jose, CA, USA). Calcein-AM was detected using a 510/20 nm bandpass (B510) filter (detection range 500-520 nm). At least 10,000 events were acquired per sample. Data were analyzed using FlowJo version 10.9. (BD Biosciences) and median fluorescence intensity (MFI) was quantified.

**Apoptosis assay.** MDA-MB-231-PacR cells expressing control (shCon) or METTL16-targeting shRNAs (shME#1 and shME#2) were treated with paclitaxel (200 nM) for 72h. Cells were harvested, washed with cold PBS, and resuspended in binding buffer. Apoptosis was measured by flow cytometry, using the Annexin V FITC Apoptosis Detection kit (BD Pharmingen, cat. # 556547) according to the manufacturer's protocol.

**Public database survival and expression analysis**

Kaplan-Meier survival analyses were performed using the online Kaplan-Meier Plotter database (http://kmplot.com), which integrates gene expression and clinical outcome data from publicly available breast cancer cohorts. METTL16 expression (probe ID: 226744_at) was analyzed in breast cancer patients. Patients were stratified into high and low expression groups using the median expression value as the cutoff, as implemented by the platform. Survival curves were generated using the Kaplan–Meier method, and statistical significance between groups was determined using the log-rank test as implemented by the platform. Hazard ratios (HRs) with 95% confidence intervals (CIs) were calculated using a Cox proportional hazards regression model provided by the database. Publicly available breast cancer transcriptomic datasets were analyzed using Breast Cancer Gene-Expression Miner v5.2 (bc-GenExMiner v5.2; http://bcgenex.ico.unicancer), an online statistical mining tool for annotated breast cancer gene-expression data (Ref). Plots comparing ABCB1 expression across TNBC subtypes were generated using the built-in expression analysis module of bc-GenExMiner v5.2. Statistical significance between groups was calculated using the Welch's t-test as implemented by the platform. For correlation analyses, pairwise gene expression correlations between ABCB1 and METTL16 were performed within the same RNA-seq cohort using the correlation module. Linear regression curves with 95% confidence intervals were generated by the platform, and correlation coefficients (r) and associated P values were calculated using Pearson's correlation test as implemented by the software.
