## Supplementary Figure Legends for "METTL16 promotes taxane resistance in Triple-Negative Breast Cancer through m^6^A-dependent translational upregulation of ABCB1"

**Suppl Figure legends:**

**Suppl Fig 1. The METTL16 gene is not amplified in PacR cells.**

**Suppl Fig 2. Taxane-resistant TNBC cells with high METTL16 are not sensitive to doxorubicin.** Cell survival assays were performed with MDAMB231 paclitaxel-sensitive and paclitaxel-resistant (PacR) cells after treatment with doxorubicin (doxo) for 48 h. ***, P< 0.001.

**Suppl Fig 3. Purity of METTL16$^{WT}$ and METTL16$^{N184A}$ used for the biochemical MTase-Glo$^{TM}$ assays. A.** B1 and B3 are the regions excised from the gel and analyzed by in-gel tryptic digest MSMS on an Orbitrap Exploris$^{TM}$ 480 Mass Spectrometer (ThermoFisher). **B.** The observed tryptic peptides from each sample are highlighted in red, confirming the identity of both proteins.

**Suppl Fig 4. Catalytic activity of METTL16 is required for paclitaxel resistance in TNBC cells. A.** Western analysis performed to determine METTL16 protein levels in MDAMB231 PacR cells overexpressing WT METTL16, N184A mutant METTL16, or empty vector. **B.** Cell survival assays were performed with the cells from **A** after treatment with paclitaxel for 3 days. The fraction of surviving cells was determined by normalizing the data from paclitaxel-treated cells to DMSO controls. **, P< 0.01; ***, P< 0.001.

**Suppl Fig 5. METTL16 regulates ABCB1 expression independently of gene amplification.** No difference in ABCB1 gene copy number **(A, B)** in PacR vs parental paclitaxel-sensitive cells, or in ABCB1 mRNA **(C, D)** in both PacR and sensitive MDAMB231 overexpressing WT METTL16 or empty vector control. **E.** ABCB1 protein level decreases in A549 cells with down-regulated METTL16**.**

**Suppl Fig 6. Verapamil increases intracellular calcein-AM accumulation in PacR TNBC cells.** WT METTL16 overexpressing MDAMB231PacR cells (WT ME-MDAMB231PacR) were pre-treated with vehicle (blue) or verapamil (5 μM, red) for 30 min, followed by incubation with Calcein-AM (0.25 μM) for 30 min. Intracellular fluorescence and the median fluorescence intensity (MFI) were analyzed by flow cytometry.

**Suppl Fig 7. METTL16 knockdown enhances paclitaxel-induced apoptosis in PacR TNBC cells.** MDAMB231 PacR cells expressing control shRNA (shCon) or METTL16-targeting shRNAs (shME-1 and shME-2) were treated with paclitaxel (200 nM) for 72 h or left untreated. Apoptosis was assessed by Annexin V-FITC and propidium iodide (PI) staining followed by flow cytometry. Live cells (Annexin V-/PI-), early apoptotic cells (Annexin V+/PI-), late apoptotic cells (Annexin V+/PI-), and necrotic cells (Annexin V-/PI+) are indicated. Percentages represent the fraction of cells within each quadrant.

**Suppl Fig 8. ABCB1 mRNA secondary structure analysis using RNAfold.** The mRNA sequence for ABCB1 (NM_001348945.2) was entered into the RNAfold Web Server. The putative METTL16-target residues (A2368, A4141, and A4497) were located in both the predicted minimum free energy and centroid secondary structures. The colors show the positional entropy value, with lower values (red) being higher confidence for the predicted structure than higher values (blue).

**Suppl Fig 9. Validation of MeRIP specificity using control primers.** MeRIP-qPCR analysis of EEF1A1 positive (Pos) and negative (Neg) control regions in MDAMB231 PacR cells expressing vector or WT METTL16. IgG was used as a negative control for the $m^6A$ immunoprecipitation. Control primers were from the Magna MeRIP $m^6A$ Kit (EMD Millipore).

**Suppl Fig 10**. **High METTL16 expression correlates with poor prognosis in TNBC.** Kaplan-Meier analysis of relapse-free survival in TNBC patients based on METTL16 expression. Patients with high METTL16 expression (red line) exhibit significantly shorter relapse-free survival compared to those with low METTL16 expression (black line).

**Suppl Fig 11. METTL16 expression positively correlates with ABCB1 in TNBC. A.** Positive correlation between METTL16 and ABCB1 in all TNBC types. **B.** High ABCB1 expression in MLIA TNBC subtype. **C.** Positive correlation between METTL16 and ABCB1 in MLIA TNBC types. All data in **A**, **B**, and **C** analyzed by Breast cancer GenEx Miner.
